# Differential usage of two, distinct DNA-binding domains regulates tissue-specific occupancy of the pioneer factor Zelda

**DOI:** 10.1101/2025.10.06.680768

**Authors:** Eliana F. Torres-Zelada, Hideyuki Komori, Hsiao-Yun Liu, Elizabeth D. Larson, Ally W.H. Yang, Zoe A. Fitzpatrick, Timothy R. Hughes, Christine A. Rushlow, Cheng-Yu Lee, Melissa M. Harrison

## Abstract

Pioneer transcription factors act at the top of gene-regulatory networks by promoting accessible chromatin at the *cis*-regulatory regions that drive gene expression. Despite their ability to bind closed chromatin, pioneer factors occupy distinct binding sites in different tissues. The pioneer factor Zelda promotes the undifferentiated fate in both the early *Drosophila* embryo and in the neural stem cells (neuroblasts) of the larval brain. Tissue-specific binding by Zelda identifies cell-type specific enhancers, which are enriched for different DNA-sequence motifs. We investigated the features that promoted cell-type specific occupancy by testing the role of conserved, structured protein domains in the capacity of Zelda to promote the embryonic and neuroblast cell fates. We unexpectedly identified that the most deeply conserved region in Zelda, the second zinc finger, has opposing functions in the embryo and neuroblasts. We showed that this zinc finger is a previously unrecognized DNA-binding domain that is specifically required for Zelda binding to a G-rich motif in larval neuroblasts. The pioneering function of Zelda depends largely on the C-terminal cluster of zinc fingers that promotes binding in the early embryo, suggesting that pioneer function may depend on how Zelda engages the genome. As opposed to co-factor expression or chromatin environment, our data identify tissue-specific usage of two, widely separated DNA-binding domains as the mechanism controlling tissue-specific binding and activity.

## Introduction

Coordinated changes in gene expression drive development of a complex organism from a single-celled zygote. Transcription factors mediate these changes in cellular identity by binding DNA and regulating specific gene-expression programs. While chromatin is a barrier to the DNA binding of many transcription factors, pioneer transcription factors bind nucleosomal DNA, establish regions of open chromatin, and facilitate the binding of additional transcription factors. In so doing, pioneer factors define the *cis*-regulatory regions that control gene expression (Iwafuchi-Doi and Zaret 2014; Zaret 2020). Because of their ability to restructure the chromatin accessibility landscape, pioneer factors act at the top of gene-regulatory networks to control developmental transitions (Larson et al. 2021b; Balsalobre and Drouin 2022; Barral and Zaret 2024; Horisawa and Suzuki 2023). Although pioneer factors bind nucleosomal DNA, their capacity to bind and open chromatin is not unrestrained. Therefore, pioneer factors both regulate development and are influenced by developmental context (Freund et al. 2024; Larson et al. 2021b). Only a subset of recognition motifs is bound *in vivo*, and binding is cell-type specific. Nonetheless, the features that regulate tissue-specific pioneer-factor occupancy remain unclear.

Because of their ability to promote broad changes in gene expression, pioneer factors drive conserved developmental transitions. One of these transitions is the initial stages of development in which the specified germ cells are reprogrammed to totipotency. At this time, the zygotic genome is transcriptionally quiescent, and maternally deposited mRNAs and proteins control the first hours of development. Only after the cells have become totipotent is widespread transcription initiated (Schulz and Harrison 2019; Vastenhouw et al. 2019). Transcriptional activation of the zygotic genome is strictly coordinated with the degradation of the maternally deposited mRNAs in what is known as the maternal-to-zygotic transition (MZT) (Vastenhouw et al. 2019). In all organisms studied to date, pioneer factors promote reprogramming of the zygotic genome and transcriptional activation (Larson et al. 2021b; Freund et al. 2024; Kobayashi and Tachibana 2021; Yartseva and Giraldez 2015). The first such pioneer factor identified was Zelda (Zld) in *Drosophila melanogaster* (Liang et al. 2008; Harrison et al. 2011; Schulz et al. 2015; Sun et al. 2015). Since its identification, Zld has served as a paradigm for understanding pioneer-factor mediated reprogramming.

In addition to its essential function in embryos, Zld also promotes the undifferentiated fate in neural stem cells (neuroblasts) of the larval brain. Neuroblasts divide asymmetrically to both self-renew and generate progeny that rapidly exits the multipotent state and begins to differentiate (Rajan et al. 2021). Failure to robustly control this transition leads to excess stem-cell divisions and tumors (Homem and Knoblich 2012; Reichardt et al. 2018). Indeed, overexpression of Zld generates extra neuroblasts and a tumor-like phenotype (Larson et al. 2021a). Despite the shared role in promoting the undifferentiated state in embryos and neuroblasts, Zld occupies distinct regions of the genome in these two cell populations (Larson et al. 2021a). In embryos, Zld binding is driven strongly by sequence with 90% of the most highly bound regions containing the canonical CAGGTA motif (Harrison et al. 2011). A cluster of four zinc fingers in the C-terminal portion of Zld mediates CAGGTA binding (Hamm et al. 2015; McDaniel et al. 2019). By contrast, Zld-bound regions in neuroblasts are only weakly enriched for a degenerate version of this motif (Larson et al. 2021a). Thus, similar to other pioneer factors, Zld has tissue-specific genome occupancy. However, the mechanisms driving the distinctive binding in embryos and neuroblasts remain unclear.

Protein-extrinsic features, like chromatin structure and cofactor expression, have been thought to be the predominant regulators of cell-type specific pioneer-factor binding (Andersson and Sandelin 2020; Heinz et al. 2015; Rada-Iglesias et al. 2011; Gibson et al. 2024; Soufi et al. 2012; Mayran et al. 2018; Buecker et al. 2014; Chronis et al. 2017; Donaghey et al. 2018). Unexpectedly, we identified a second, previously unappreciated DNA-binding domain in the paradigmatic pioneer factor Zld and demonstrate that differential usage of the two widely separated DNA-binding domains governs the tissue-specific binding and activity of Zld. We show that the second zinc finger in Zld, which is the most deeply conserved portion of the protein, is required for binding *cis*-regulatory regions in neuroblasts, but not in the embryo. Furthermore, this differential usage of these two DNA-binding domains regulates the pioneering function of Zld, which largely depends on the C-terminal cluster of zinc fingers that serves as the embryonic DNA-binding domain. Our studies show that developmental context can control pioneer activity through regulation of DNA-binding domain usage and suggest that, in addition to protein-extrinsic features, protein-intrinsic regulation control tissue-specific, pioneer-factor binding.

## Results

### The deeply conserved second zinc finger in Zelda is required to promote the undifferentiated neuroblast fate

Zld is a large protein, comprised predominantly of unstructured regions with six, structured C2H2 zinc fingers (Fig. 1a). Prior work identified the C-terminal cluster of four zinc fingers as the DNA-binding domain required for activating embryonic Zld-target genes (Hamm et al. 2015). The second zinc finger (ZnF2) is the most deeply conserved portion of the protein (Ribeiro et al. 2017), and we previously showed that this domain inhibits Zld-mediated activation of zygotic transcription (Hamm et al. 2017). Because Zld promotes the undifferentiated fate in both the early embryo and neuroblasts in the larval brain (Liang et al. 2008; Larson et al. 2021a), we tested the roles of the conserved, structured zinc-finger domains of Zld in promoting the pluripotent neuroblast fate. We generated transgenic lines overexpressing Zld variants with targeted mutations in individual zinc fingers using a heat-inducible, pan-neuroblast driver (*Wor-Gal4*, *Tub-Gal80^ts^)* and quantified type II neuroblasts in third instar larval brain lobes. Because there are always exactly eight type II neuroblasts per lobe in the larval brain, extra neuroblasts are indicative of changes to cell fate. Similar to our previous work, we demonstrated that overexpression of Zld resulted in extra neuroblasts and that mutation to the cluster of four DNA-binding zinc fingers (ZnF3-6) partially impaired this ability of Zld to promote neural stem cell identity (Fig. 1b; Larson et al. 2021.) As predicted from the endogenous mutants (Hamm et al. 2017), mutations in the first zinc finger (ZnF1) resulted in extra neuroblasts, similar to wild-type protein expression (Fig. 1b). Based on our prior demonstration of the inhibitory function of ZnF2 (Hamm et al. 2017), we predicted that mutations in this domain would further enhance the ability of Zld expression to drive supernumerary neuroblasts. In contrast to this prediction, mutations in ZnF2 completely abrogated Zld-mediated neuroblast induction (Fig. 1b). To test whether this counterintuitive result reflected the endogenous function of Zld, we tested whether mutations to ZnF2 phenocopied *zld* mutants in the larval brain. Although Zld is not required for neuroblast formation, null alleles in *zld* suppress the tumor-like phenotype caused by mutations in *brain tumor* (*brat*) (Larson et al. 2021a; Reichardt et al. 2018). Phenocopying null alleles, males hemizygous for a mutation in ZnF2 suppress the extra neuroblasts caused by mutations in *brat* (Fig. 1c). Together these data reveal that the function of ZnF2 is context-dependent: required to promote the neuroblast fate, but inhibiting the role of Zld in embryonic gene expression.

**Figure 1:**
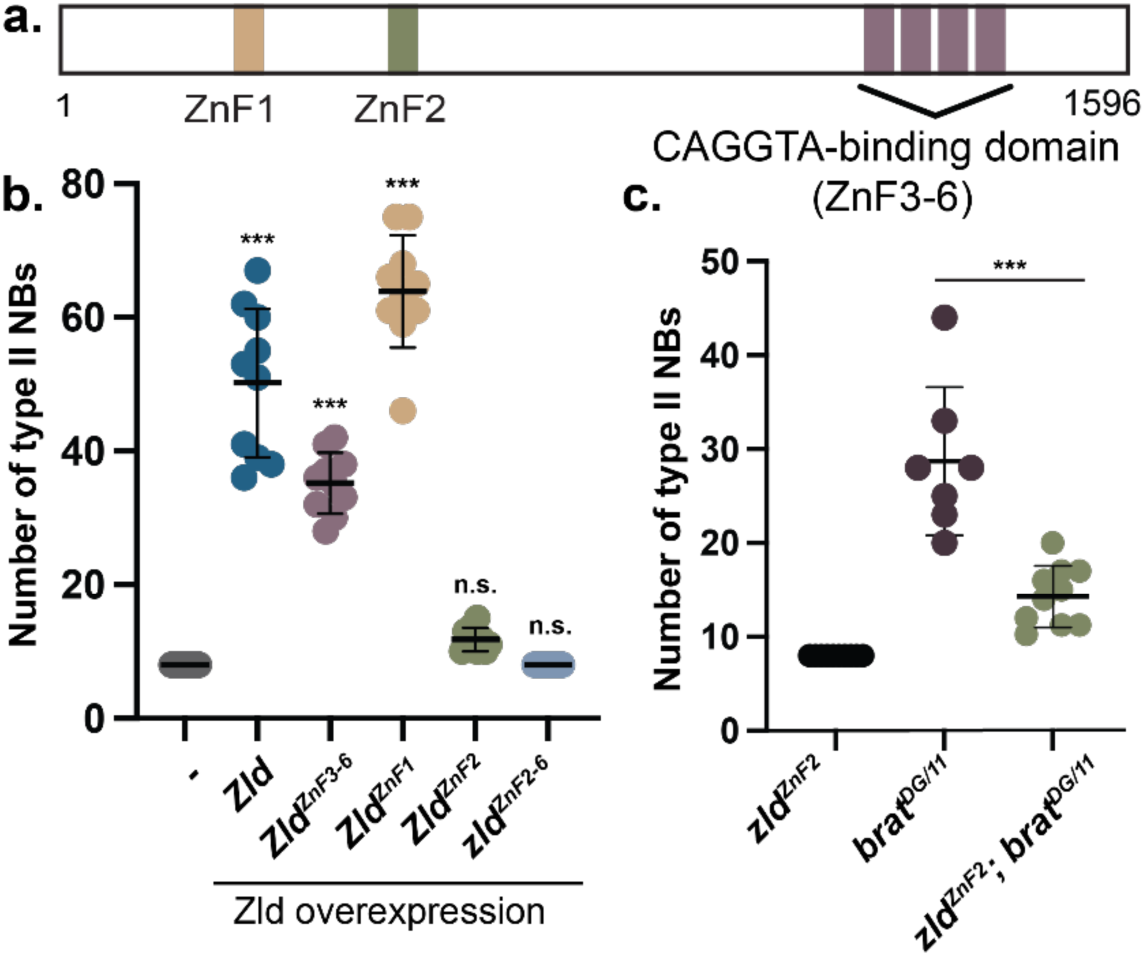
The second zinc finger of Zld (ZnF2) is required to promote the neural stem cell fate. **a.** Schematic of Zld including the six zinc fingers as indicated. **b.** Number of type II neural stem cells per brain lobe. The indicated Zld constructs were driven in stem cells by *Wor-Gal4*, *Tub-Gal80^ts^*. Comparisons are to *Wor-Gal4*, *Tub-Gal80^ts^* alone using an ANOVA. **c.** Number of type II neural stem cells per brain lobe in the genotypes indicated. Males were used for all assays. For Zld^ZnF2^, the single copy of *zld* on the X chromosome carried the mutation. Genotypes were compared using an unpaired t-test. For each genotype: n = 10 brains, ***, p < 0.001. ns = not significant.

### Differential contributions of ZnF2 and ZnF3-6 to Zld-motif recognition and target-site selection

Having identified tissue-specific requirements for ZnF2, we used cell culture to determine if these differences were due to the cellular context. Exogenous expression of Zld in S2 cells robustly activates transcription driven by the *scute* promoter, an established embryonic Zld-target gene containing four canonical CAGGTA motifs in the promoter (Fig. 2a) (Hamm et al., 2015). As we previously demonstrated, expression of Zld with mutations in ZnF2 (Zld^ZnF2^) significantly hyperactivated the *scute* reporter as compared to wild type, while Zld^ZnF3-6^ reduced activation (Fig. 2a) (Hamm et al. 2015, 2017). We generated a reporter for Zld activity in neuroblasts by cloning a Zld-bound, neuroblast-specific enhancer of the type II neuroblast master regulator *tailless* (*tll*) upstream of the Drosophila Synthetic Core Promoter (DSCP) driving firefly luciferase. Like most neuroblast-specific, Zld-target genes, the *tll* enhancer lacks the canonical CAGGTA Zld-binding motif that defines embryonic target genes (Harrison et al. 2011; Larson et al. 2021a; Nien et al. 2011). We confirmed that Zld expression activates the *tll* enhancer in S2 cells (Fig. 2b). In contrast to the *scute* reporter, Zld-mediated activation depended on ZnF2, but not ZnF3- 6 (Fig. 2b), reflecting the opposing activities observed for ZnF2 *in vivo*. These data indicate that the activity of ZnF2 is not controlled by the cellular environment, but rather by the underlying DNA-regulatory element.

**Figure 2:**
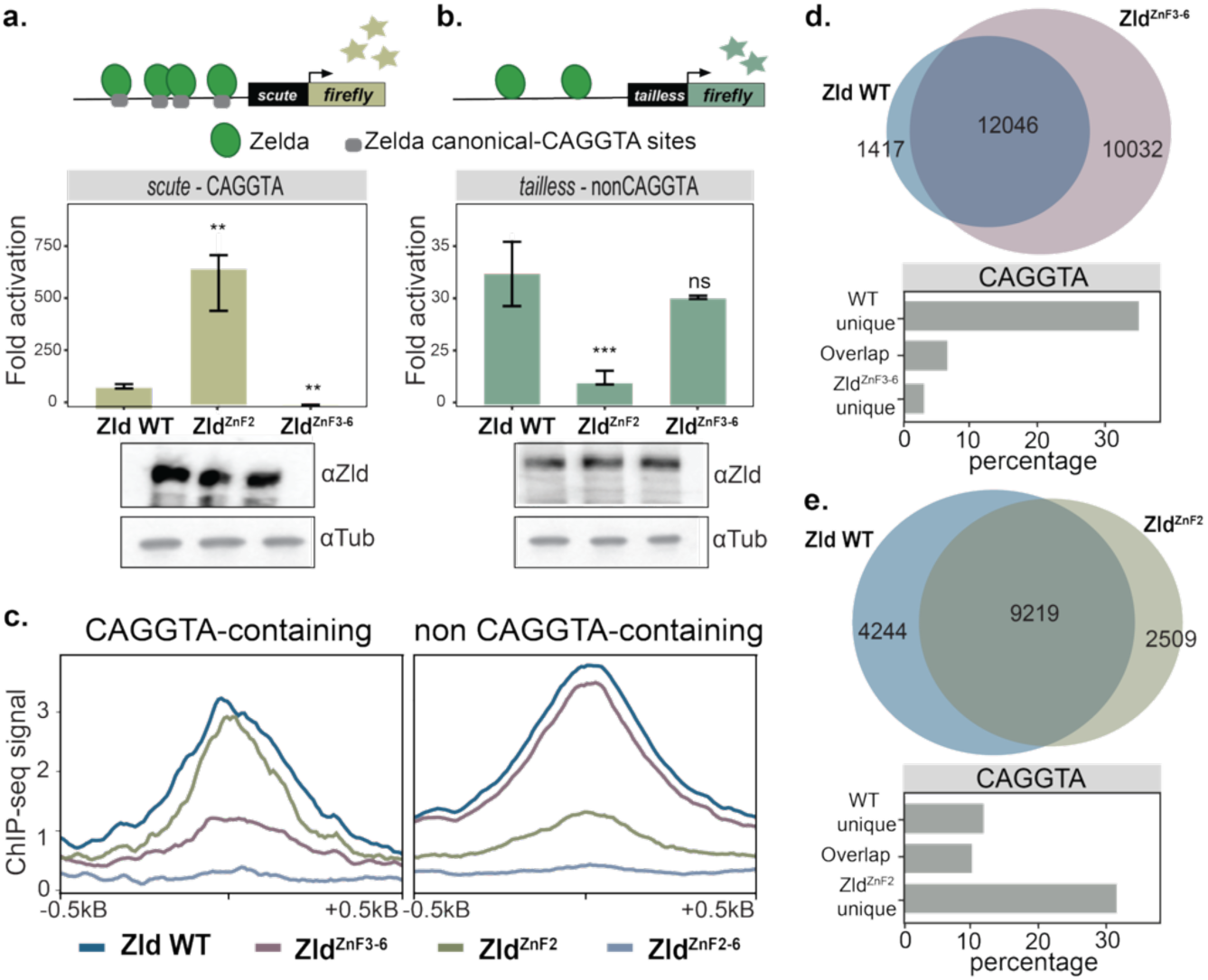
Tissue-specific activity of ZnF2 is differentially interpreted based on the underlying DNA-sequence. **a-b.** Representative luciferase reporter assays of Zld activity for wild-type (WT), ZnF2-mutant and ZnF3-6-mutants (*top*). Each mutated version is compared to Zld WT using a t-test. n = 3 technical replicates. **, p < 0.01, ***, p < 0.001. ns = not significant. Immunoblots for Zld and tubulin, demonstrating similar expression levels (*bottom*). The s*cute* promoter used in **a.** contains four CAGGTA elements. The *tailless* reporter used in **b.** is driven by the previously defined neuroblast-specific enhancer. **c.** Metaplots of Zld ChIP-seq in S2 cells expressing the Zld proteins indicated below. Signal is shown for binding to regions of closed chromatin containing CAGGTA sites (*left*) or not containing CAGGTA sites (*right*). **d.** Venn diagram of all ChIP-seq peaks for wild-type Zld and Zld^ZnF3-6^ (*above*). Percentage of peaks that contain the canonical Zld-binding motif, CAGGTA, in each category (*below*). **e.** Venn diagram of all ChIP-seq peaks for wild-type Zld and Zld^ZnF2^ (*above*). Percentage of peaks that contain the canonical Zld-binding motif, CAGGTA, in each category (*below)*.

To more directly investigate the functions of ZnF2 and ZnF3-6, we sought to assay how mutations to these domains influenced chromatin binding. For this purpose, we generated stable S2 cell lines allowing for inducible expression of wild-type Zld or Zld variants harboring mutations in ZnF2, ZnF3-6 or both domains (Zld^ZnF2^, Zld^ZnF3-6^, Zld^ZnF2-6^) and used chromatin immunoprecipitation coupled with high-throughput sequencing (ChIP-seq) to assay binding. Zld binds promiscuously to accessible chromatin (Gibson et al. 2024). Therefore, we focused our initial analysis on regions at which Zld functions as a pioneer factor; binding to closed chromatin and promoting accessibility (Gibson et al. 2024). We subdivided these Zld-pioneered sites depending on whether they contained a canonical CAGGTA Zld-binding motif (Fig. 2c). As expected, Zld binding was lost at all Zld-pioneered sites when ZnF2-6 were mutated (Fig. 2c). Mutations in the previously defined DNA-binding domain (ZnF3-6) resulted in a loss of chromatin binding at regions containing a CAGGTA motif, but Zld retained binding at loci lacking this canonical motif (Fig. 2c). By contrast, ZnF2 was dispensable for binding CAGGTA-motif containing regions but required for binding other loci (Fig. 2c). We extended our analysis to include all ChIP-seq peaks and looked at the overlap between wild-type Zld occupancy and that of proteins with mutations in ZnF2 or ZnF3-6 (Fig. 2d-e). Mutation in ZnF3-6 resulted in a loss of Zld binding at sites enriched for the CAGGTA motif (WT unique, Fig. 2d.) When ZnF2 was mutated, sites that gained binding were enriched for the CAGGTA motif, suggesting that ZnF2 activity displaces Zld from CAGGTA motifs (ZnF2 unique, Fig. 2e). These results suggest that ZnF2 and ZnF3-6 mediate distinct and independent activities that regulate DNA binding with ZnF3-6 promoting binding to CAGGTA elements and ZnF2 promoting binding to noncanonical elements.

Our data suggested that ZnF2 promotes binding of Zld to non-CAGGTA-containing regions. Zld-bound, neuroblast-specific regulatory elements are largely devoid of CAGGTA motifs. Instead, they are enriched for a GA-dinucleotide motif bound by several factors, including the pioneer factor GAGA factor (GAF) (Larson et al. 2021a). This enrichment led us to query whether GAF might promote Zld binding via ZnF2. In the early embryo, GAF and Zld occupy many overlapping sites, but this binding is largely independent (Gaskill et al. 2021). Nonetheless, a recent study identified a redistribution of Zld binding in the early embryo upon mutation to ZnF5, which is essential for the CAGGTA-binding activity of Zld, and suggested that this redistribution may be driven by GAF (Fallacaro et al. 2024). We therefore further investigated the relationship between Zld and GAF binding using our cell-culture system. We did not identify a specific enrichment of the GAF-binding GA-dinucleotide motif in regions that were specifically gained or lost upon mutations in ZnF2 or ZnF3-6 (Supp. Fig. 1a-b). Furthermore, we depleted GAF using RNAi, confirmed that Zld levels remained unchanged and identified Zld-binding sites using ChIP-seq (Supp. Fig. 1c-f). Like our results in the embryo (Gaskill et al. 2021), Zld binding is retained upon GAF depletion at both sites driven by ZnF2 and ZnF3-6 (Supp. Fig. 1d-f), suggesting that GAF is not broadly regulating Zld occupancy.

### The DNA-binding activity of Zld is mediated by two distinct, structured domains with unique motif preferences

Together our analyses support a role for ZnF2 in promoting Zld chromatin occupancy at the non- CAGGTA-containing loci essential for promoting neuroblast fate. Nonetheless, it remained unclear whether ZnF2 promoted occupancy directly through interactions with DNA or indirectly. To address this, we expressed and purified recombinant Zld constructs containing either the N-terminus, including ZnF2, (Zld N) or the C-terminus, including ZnF3–6, (Zld C) and assessed their binding specificity using protein binding microarrays (PBMs) (Fig. 3a) (Weirauch et al. 2014). As expected, Zld C preferentially binds the canonical CAGGTA motif (Fig. 3a) (Brennan et al. 2023). By contrast, Zld N showed the highest affinity for a G-rich variant motif (AAGGGT) (Fig. 3a). To test the affinity of Zld N for the newly identified G-rich motif, we performed electrophoretic mobility shift assays (EMSAs) using a CAGGTA-containing probe derived from the *scute* promoter that has been used previously (Hamm et al. 2015; McDaniel et al. 2019; Harrison et al. 2011). To assess binding specificity, we generated a variant of this probe in which the CAGGTAG site was replaced with the identified AAGGGT motif. Zld C specifically binds to the CAGGTAG-containing probe, while Zld N preferentially binds to the AAGGGT variant (Fig. 3b). Neither protein bound to a probe in which this sequence was mutated (Fig. 3b). Binding by Zld N to the AAGGGT- containing probe depended largely on ZnF2 as mutation severely impaired the ability of the protein to shift the probe (Fig. 3c-d). These *in vitro* studies strongly support a direct interaction between ZnF2 and the AAGGGT motif.

**Figure 3:**
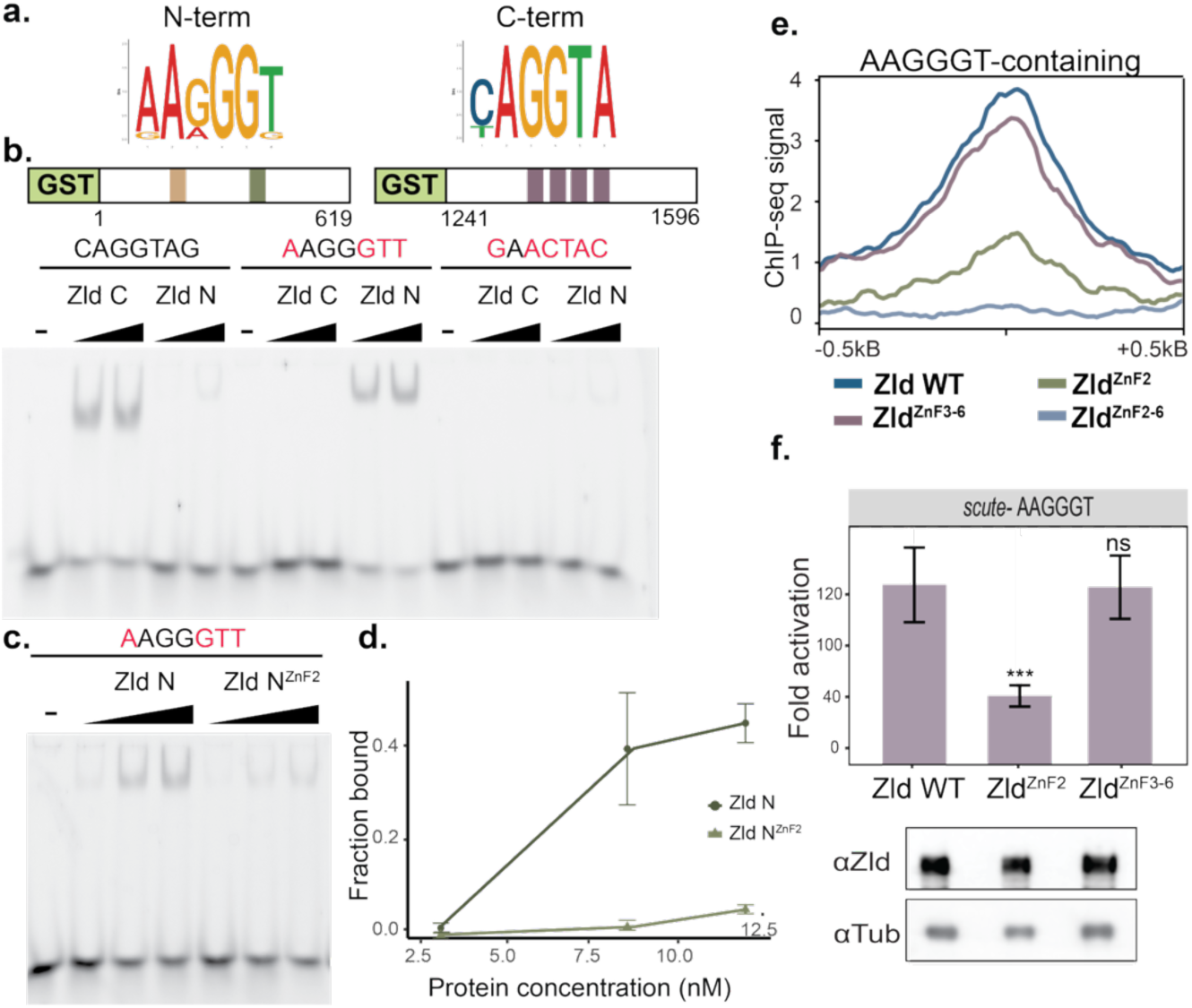
Zelda contains two, widely separated DNA-binding domains. **a.** Position-weight matrices from protein-binding microarrays based on binding by recombinant Zld constructs. Zld N is an N-terminal construct containing ZnF2. Zld C is a C-terminal construct containing ZnF3-6. **b.** Models of the Zld constructs used for EMSA (*top*). Representative EMSA showing binding of Zld N or Zld C to CAGGTAG, AAGGGTT, or a mutated motif. **c.** Representative EMSA of AAGGGTT-containing sequence with wild-type Zld N or Zld N with a mutated ZnF2. **d.** Quantification of the fraction of DNA bound (y-axis) as a function of protein concentration (x-axis) for two replicate EMSAs as in **c**. Error bars are ± standard deviation from n = 2 replicates. **e.** Metaplots of Zld ChIP-seq in S2 cells expressing the Zld proteins indicated below. Signal is shown for binding to regions of closed chromatin containing AAGGGT motifs. **f.** Representative luciferase assays as in Figure 2a in which the CAGGTA motifs in the *scute* promoter were replaced with AAGGGT. Each Zld-mutated version is compared to Zld WT using t-test. n = 3 technical replicates. ***, p < 0.001. ns = not significant. Immunoblots for Zld and tubulin, demonstrating similar expression levels (*bottom*).

To investigate the role of the AAGGGT motif in directing Zld binding in cells, we determined how Zld occupancy at this motif was affected by mutations in ZnF2 or ZnF3-6. We showed that binding to AAGGGT motifs in closed chromatin depended on Zld^ZnF2^, but not Zld^ZnF3-6^, when compared to wild type (Fig. 3e). Thus, in both culture and *in vitro* ZnF2 promotes binding to AAGGGT motif-containing DNA. We further tested the functional significance of this binding by replacing the CAGGTAG sites in the *scute* luciferase reporter with the G-rich motif and testing for activation by Zld, Zld^ZnF2^ and Zld^ZnF3-6^. Substitution of the four CAGGTAG sites converted the *scute* reporter from depending on ZnF3-6 for activation to depending on ZnF2 (Fig. 3f), essentially making this reporter mimic the neuroblast-specific *tll* reporter. Thus, ZnF2 is a previously unidentified DNA-binding domain in the paradigmatic pioneer factor Zld.

### ZnF2 regulates context-specific Zelda binding and is essential for promoting an undifferentiated fate in neuroblasts

Our identification of ZnF2 as a newly identified DNA-binding domain specifically required in neuroblasts suggested that differential usage of the two DNA-binding domains in Zld could drive the tissue-specific occupancy. To test whether ZnF2 is broadly required for Zld chromatin occupancy in neuroblasts, we leveraged our transgenic system to express Zld and mutated versions of Zld in *brat-*mutant brains to enrich for type II neuroblasts, as we did previously (Larson et al. 2021a). Our prior data showed that mutations in *brat* result in thousands of type II neuroblasts that resemble endogenous neuroblasts in their gene expression profiles as well as their ability to respond to developmental cues and differentiate (Rives-Quinto et al. 2020; Lee et al. 2006). These and other data support our use of the *brat* mutant to enrich for type II neuroblasts, providing a powerful system for genomic analysis (Larson et al. 2021a; Rives-Quinto et al. 2020; Shen et al. 2025). While the most straightforward experiment would be to perform analysis on animals in which ZnF2 is endogenously mutated, these are not feasible as mutations to this domain, like knockdown of *zld* by RNAi, suppress the supernumerary neuroblast phenotype, changing the cell-type composition of the brain (Fig. 1c) (Reichardt et al. 2018). We therefore expressed HA-tagged versions of Zld that were either wild-type or mutated for ZnF2 or ZnF3-6 and performed anti-HA CUT&RUN to identify Zld-bound loci. The HA-tag did not inhibit Zld function, as expression in neuroblasts resulted in extra neural stem cells, similar to induced expression of untagged Zld (Supp. Fig 2a-c). To control for the fact that these proteins are being exogenously expressed, we compared all binding sites to exogenously expressed wild-type Zld. Exogenously expressed wild-type Zld binds to largely the same loci as endogenous Zld using antibodies recognizing both Zld and HA for CUT&RUN (Fig. 4a, Supp. Fig 2d-e). Overexpression results in additional binding sites but comparing between transgenically expressed wild-type and mutant protein allowed us to control for these novel binding sites (Fig. 4a, Supp. Fig 2d). Our analysis demonstrated that ZnF2, but not ZnF3-6 was broadly required for Zld occupancy, including the neuroblast-specific *tll* enhancer (Fig. 4a,b). Notably, ZnF2-dependent sites were enriched for the AAGGGT motif (Fig. 4c).

**Figure 4:**
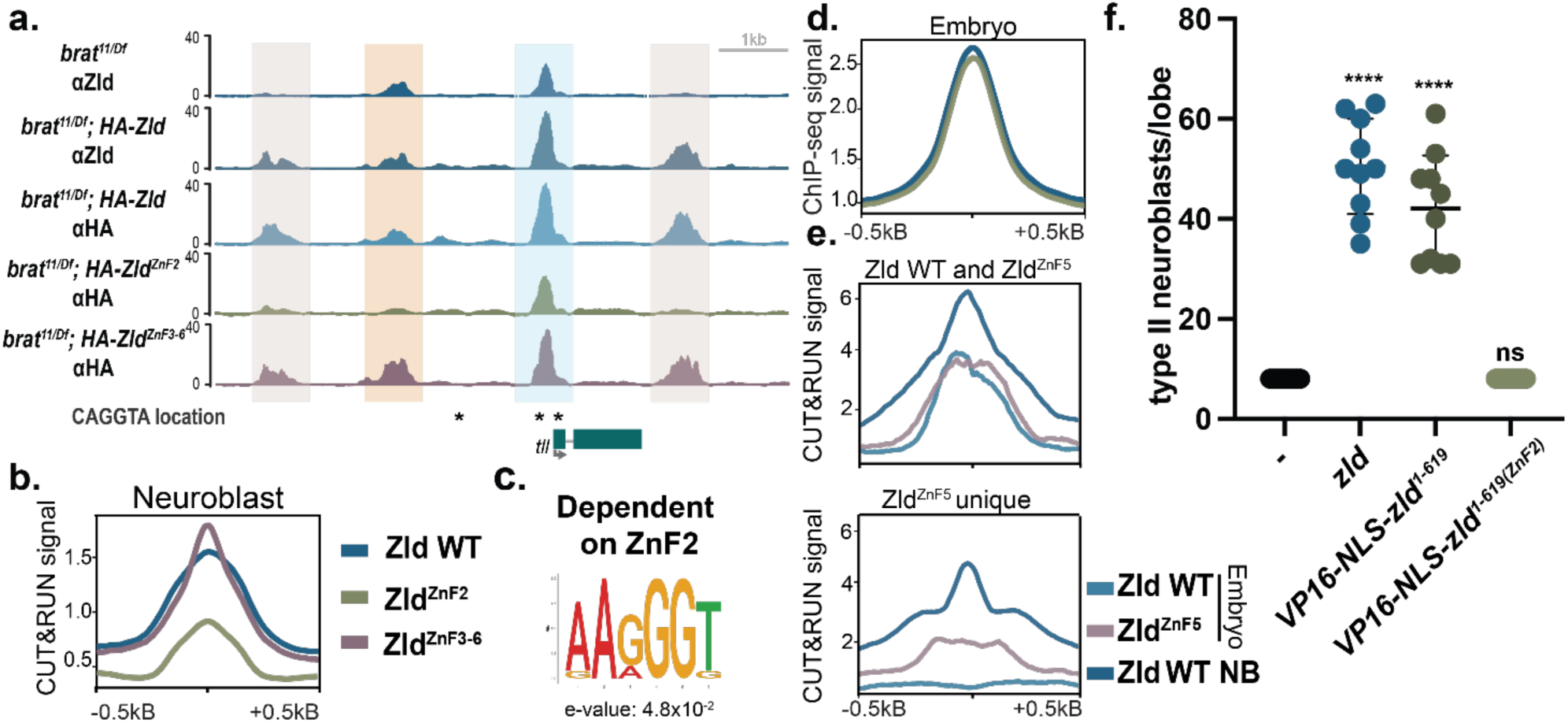
The N-terminus of Zld is necessary and sufficient for promoting the stem-cell fate. **a.** Genome browser tracks of CUT&RUN for HA or Zld, as indicated, and centered on the *tll* locus. Experiments were performed on *brat-*mutant brains to enrich for type II neuroblasts. Highlighted regions: Orange, the neuroblast-specific enhancer that requires ZnF2, but not ZnF3-6 for Zld binding. Blue, the promoter that is occupied in all conditions. Gray, two regions that are bound by Zld only upon overexpression. These also depend on ZnF2 for occupancy but can be identified in comparison to CUT&RUN on endogenously expressed Zld. **b.** Metaplot of average z-score normalized anti-HA CUT&RUN signal from *brat* larval brains for the genotypes indicated. **c.** Motif enrichment for Zld-bound sites that are dependent on ZnF2 in neuroblasts (determined by MEME-suite). **d.** Metaplots of spike-in normalized ChIP-seq for Zld from either wild-type or Zld^ZnF2^ embryos. **e.** Metaplots of z-score normalized CUT&RUN signal on endogenous Zld and Zld^ZnF5^ in stage 5 embryos (from Fallacaro et al. bioRxiv 2024) and wild-type Zld from neuroblasts. Regions bound by both wild-type Zld and Zld^ZnF5^ (*above*) and regions unique to Zld^ZnF5^ (*below*). **f.** Number of type II neuroblasts per brain lobe in which the indicated VP16-NLS Zld constructs are expressed. Comparisons are to *Wor-Gal4*, *Ase-Gal80* (-) alone using a one-way ANOVA. For each genotype: n = 10 brains, ****, p < 0.0001. ns = not significant.

Our data identify ZnF2 as a DNA-binding domain essential for the binding profile of Zld in the larval neuroblasts. In contrast to the neuroblasts, the CAGGTA sequence is a major driver of Zld occupancy in embryos (Harrison et al. 2011; Larson et al. 2021a; Nien et al. 2011). This suggests that recognition of the AAGGGT motif by ZnF2 is unlikely to drive Zld binding at this early stage of development, and its activity might impede the recruitment of Zld to embryo-specific *cis*-regulatory elements. We performed spike-in normalized ChIP-seq for Zld in wild-type embryos (*w^1118^*) or embryos laid by mothers homozygous for *zld^ZnF2^* (Hamm et al. 2017). Importantly, Zld is expressed at equivalent levels in control (*w^1118^*) and *zld^ZnF2^* embryos (Supp. Fig. 3a). Mutation of ZnF2 did not significantly change the genome occupancy of Zld in the embryo (Fig. 4c, Supp. Fig. 3b), indicating that ZnF2 is selectively required for Zld binding in neuroblasts but dispensable for its genome-wide binding in the embryo. Previously identified Zld-target genes that were upregulated in the embryo when ZnF2 is mutated did not show corresponding changes in Zld binding, suggesting that transcriptional effects may not arise from differential occupancy (Supp. Fig 3c) (Hamm et al. 2017). We then leveraged CUT&RUN data from embryos in which ZnF5 was mutated, abrogating binding to the canonical CAGGTA motif (Fallacaro et al. 2024). We analyzed these data to determine if ZnF2 might promote binding to neuroblast-specific regulatory elements in the embryo when CAGGTA binding was disrupted. We compared the genome-wide binding profiles and identified loci shared between wild-type (WT) and Zld^ZnF5^ mutant embryos, as well as those uniquely bound in either condition. Sites specifically bound in the ZnF5 mutant might represent new binding events driven by ZnF2. Supporting this model, regions bound by Zld^ZnF5^ (both unique and shared with wild type) were strongly bound by Zld in neuroblasts (Fig. 4e, Supp. Fig. 3d). Consistent with a role for ZnF2 in mediating this redistribution, Zld binding at ZnF5-unique sites was reduced in neuroblasts expressing the ZnF2 mutant (Supp. Fig. 3d).

Finally, we asked whether the N-terminal portion of Zelda (aa 1-619), which includes ZnF2 but lacks ZnF3-6, was sufficient to activate Zld-responsive loci and promote neuroblast fate *in vivo*. Because this fragment lacks the endogenous activation domain (aa 904–1297) (Hamm et al. 2015), we fused it to the potent VP16 transcriptional activation domain (Janssens et al. 2014). To ensure proper localization, a nuclear localization signal (NLS) was included (Supp. Fig. 4a). Expression of VP16-NLS-Zld^1–619^ in type II neuroblasts driven by *Wor-Gal4, Tub-Gal80^ts^* resulted in extra neuroblasts, comparable to expression of the full-length protein (Fig. 4f, Supp. Fig. 4b). Importantly, this effect was abolished when ZnF2 was mutated, indicating that ZnF2 is required for this activity (Fig. 4f). Further supporting the specificity of this effect, VP16-NLS-Zld^1117-1487^, which includes ZnF3-6 but not ZnF2, failed to induce extra neuroblasts. To test whether the ability to promote extra neuroblasts likely reflected Zld-mediated transcriptional activation, we determined whether VP16-NLS-Zld^1–619^ could promote expression of our neuroblast-specific reporters in tissue culture. Indeed, VP16-NLS-Zld^1–619^ activated both the *scute* reporter with AAGGGT motifs and the *tll* reporter. By contrast, mutation of ZnF2 in this construct failed to activate despite comparable protein levels (Supp. Fig. 4c). Together, these results demonstrate that the N-terminal region of Zld, including ZnF2, is sufficient to confer neuroblast-specific activity.

### Zld pioneering activity depends largely on ZnF3-6

While pioneer factors bind nucleosomal DNA and promote local chromatin opening, most pioneer factors only promote chromatin accessibility at a subset of bound regions (Gibson et al. 2024; Soufi et al. 2012; Veil et al. 2019). Thus, in many cases pioneer binding is separable from activity (Freund et al. 2024). We therefore investigated how ZnF2 and ZnF3-6 contribute to the capacity of Zld to promote chromatin accessibility. We performed assays for transposase-accessible chromatin coupled with high-throughput sequencing (ATAC-seq) in cells expressing wild-type Zld or Zld with mutations in ZnF2 or ZnF3-6, similar to our ChIP-seq to assay binding. We previously showed that Zld promotes chromatin accessibility at a subset of bound regions (Gibson et al. 2024). Mutation of ZnF2 abrogated binding and the ability of Zld to generate chromatin accessibility at these regions (Fig. 5a). Even though Zld^ZnF3-6^ largely retained binding at these regions, this binding did not promote chromatin accessibility (Fig. 5a). Thus, at these regions ZnF2 is required for pioneer-factor binding but ZnF3-6 is needed for pioneering activity. When we analyzed the effects of these ZnF mutants genome wide, we found that mutations in ZnF3-6 had a significantly more pronounced effect on chromatin accessibility than mutation of ZnF2 despite the broader requirement for ZnF2 in DNA binding (Fig. 5b). At regions that contain the ZnF2 binding motif, AAGGGT, mutation of ZnF3-6 has an equivalent effect on chromatin accessibility as does mutation in ZnF2, which disrupts binding (Fig. 5b). These results demonstrate that Zld pioneering activity depends largely on the previously identified, canonical DNA-binding domain, ZnF3-6, regardless of whether this domain drives DNA binding.

**Figure 5:**
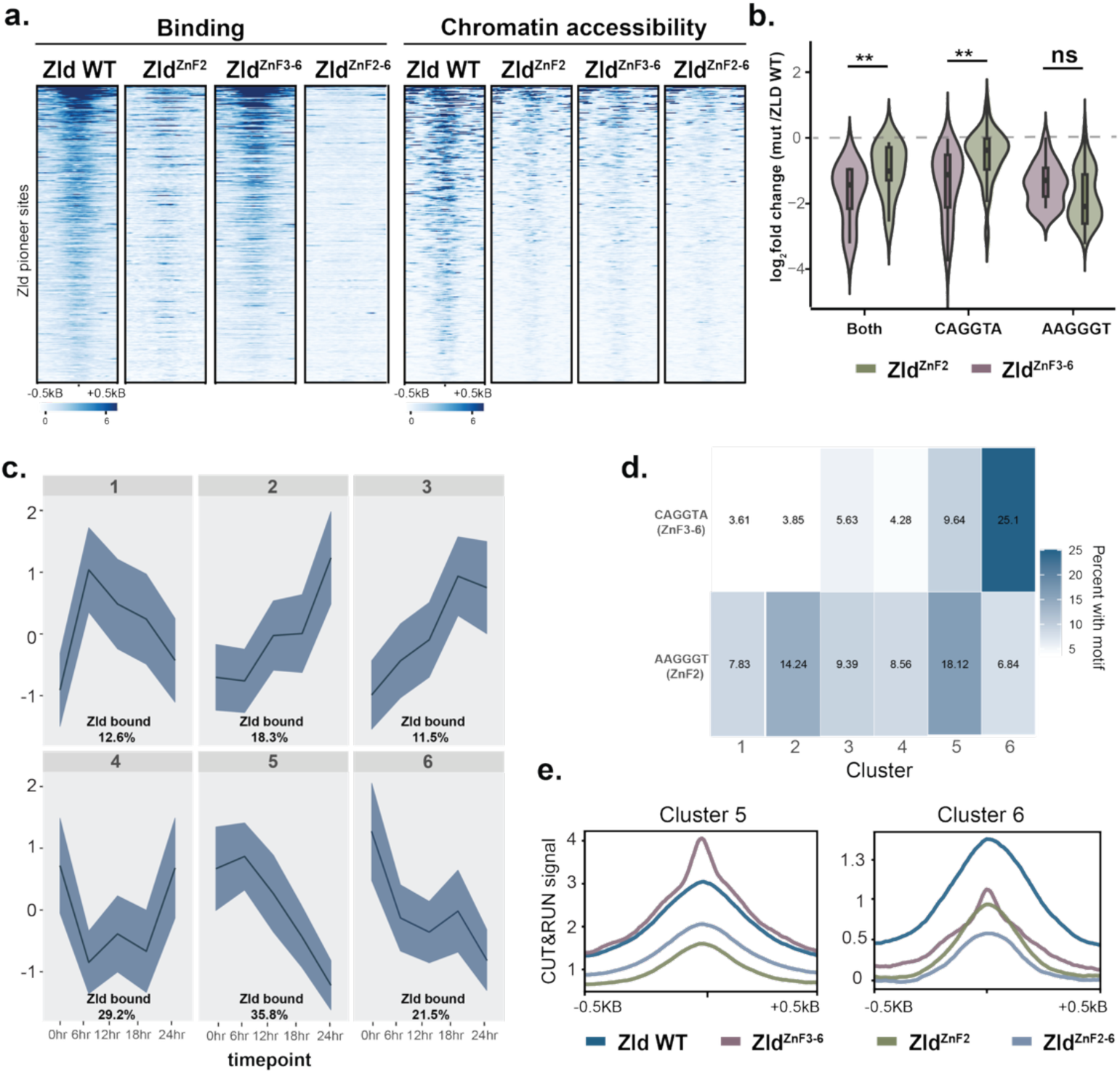
The pioneering activity of Zld depends largely on the C-terminal cluster of zinc fingers. **a.** Heatmaps of z-score normalized Zld ChIP-seq signal for Zld WT and indicated mutants at loci where wild-type Zld expression in S2 cells promotes chromatin accessibility (*left*). ATAC-seq from the same cell populations (*right*). Heatmaps are centered on peak summits (±500 bp). **b.** Violin plot of log₂ fold change in ATAC-seq signal (Zld^ZnF3-6^ or Zld^ZnF2^ over Zld WT) at Zld-pioneered sites grouped by underlying DNA motifs. Non-parametric Wilcoxon test **, p < 0.01. ns = not significant. **c.** k-means clustering of regions that change in accessibility based on ATAC-seq during induced, synchronous type II neuroblast differentiation (from Larson et al. 2021a). Percent of Zld-bound sites in each cluster is indicated. Regions in class 5 and 6 decrease in chromatin accessibility but with different dynamics. **d.** Percent of regions in each ATAC-seq cluster that contains the indicated Zld-binding motif. **e.** Metaplots of average z-score normalized anti-HA CUT&RUN signal from *brat*-mutants brains expressing the indicated HA-Zld transgenes. Metaplots are centered on regions of dynamic ATAC-seq accessibility.

Zld is a well-defined pioneer factor in the early embryo (Brennan et al. 2023): Zld binds to nucleosomes *in* vitro, is required for chromatin accessibility, and facilitates transcription-factor binding and gene expression (Foo et al. 2014; Liang et al. 2008; Xu et al. 2014; Yáñez-Cuna et al. 2012; Schulz et al. 2015; Sun et al. 2015; McDaniel et al. 2019). Nonetheless, the relationship between Zld, chromatin accessibility and gene expression remains less clear in the neuroblasts. We previously used k-means clustering to define six classes of loci based on their changes in chromatin accessibility in a system that mimic *in vivo* neuroblast differentiation (Fig. 5c) (Larson et al. 2021a; Rives-Quinto et al. 2020). Our analysis showed that cluster 6, which contained regions that rapidly lost accessibility following the exit from the stem-cell fate, was enriched for the CAGGTA motif bound by ZnF3-6 and suggested that Zld may function as a pioneer factor at sites containing this motif (Larson et al. 2021a). Indeed, Zld-bound regions with the CAGGTA motif lose accessibility over the time course while Zld-bound regions lacking this motif remain accessible. We reanalyzed these data for enrichment of the newly identified ZnF2- binding motif, AAGGGT, focusing on clusters 5 and 6 that are enriched for Zld-binding sites and lose chromatin accessibility over differentiation. In contrast to cluster 6 that is enriched for the CAGGTA motif, cluster 5 shows the strongest enrichment for the AAGGGT motif (Fig. 5d). When we looked at how ZnF2 and ZnF3-6 differentially contribute to Zld binding at these regions, we identified that ZnF3-6 promoted binding to regions in cluster 6, but not cluster 5, and that mutation in ZnF2 had a stronger effect on Zld binding at regions in cluster 5 (Fig. 5e). These data support a model in which the pioneering activity of Zld is driven by the C-terminal cluster of zinc fingers (ZnF3-6) and that this pioneer activity may not be required at many Zld-bound regions in the neuroblasts where binding is mediated by ZnF2.

## Discussion

The ability of pioneer factors to access their binding motifs in the context of closed chromatin enables them to act at the top of gene-regulatory networks to drive widespread changes in cell identity (Larson et al. 2021a; Balsalobre and Drouin 2022; Barral and Zaret 2024; Horisawa and Suzuki 2023; Lee et al. 2005, 2019). Nonetheless, these factors show tissue-specific chromatin occupancy. Chromatin structure, co-factor expression and local protein concentration all shape tissue-specific pioneer factor binding (Gibson et al. 2024; Freund et al. 2024; Soufi et al. 2012; Mayran et al. 2018). In our investigation of the paradigmatic pioneer factor Zld, we unexpectedly identified a second, previously uncharacterized DNA-binding domain and showed that preferential usage of the two, widely separated DNA-binding domains governs the distinct Zld-binding profiles identified in the early embryo and neuroblasts. Furthermore, the two well-structured domains differentially contribute to the pioneering activity of Zld, suggesting that Zld may use disparate mechanisms to activate transcription in neuroblasts and the embryo.

Our studies were initiated by the identification of opposing functions for the deeply conserved second zinc finger of Zld in the embryo and neuroblasts: ZnF2 is required for promoting neuroblast fate but inhibits Zld-mediated transcriptional activation in the embryo (Hamm et al. 2017). Our demonstration that ZnF2 encodes a neuroblast-specific DNA-binding domain explains these results. ZnF2 is required for Zld binding to the *cis*-regulatory elements that drive neuroblast fate. These regulatory elements lack the canonical CAGGTA motif bound by Zld in embryos and are instead enriched for the G-rich motif bound by ZnF2. By contrast, the *cis*-regulatory elements that promote zygotic genome activation are highly enriched for the CAGGTA motif bound by ZnF3-6 (Harrison et al. 2011; Nien et al. 2011). Indeed, the CAGGTA motif is the most enriched element underlying binding of the transcription factors that pattern the early embryo, even more abundant than the motif bound by the assayed factor (Li et al. 2008; Nien et al. 2011). In addition, the abundance of CAGGTA motifs are predictive of transcriptional activity (Bosch et al. 2006). We propose that ZnF2 inhibits Zld activity in the embryo by competing with ZnF3-6 occupancy of these CAGGTA elements. While we did not identify an enrichment of Zld occupancy at CAGGTA motifs in embryos with ZnF2 mutated, our studies in cell culture showed that mutations in ZnF2 promoted occupancy at regions containing this sequence. The ChIP-seq data suggest that mutation of ZnF2 does not affect steady-state levels of Zld binding at specific elements. Zld interactions with chromatin are highly dynamic, and we propose that mutations to ZnF2 might stabilize these dynamic interactions at CAGGTA elements.

Recent studies of transcription factors, including Zld, have largely focused on the role of intrinsically disordered regions in promoting chromatin occupancy (Ferrie et al. 2022; Kumar et al. 2023; Fallacaro et al. 2024). In contrast to models that focus on low-affinity interactions mediated by disordered domains in governing tissue-specific binding, our data point to the role of a structured, conserved domain. In embryos mutation of ZnF5, which disrupts the CAGGTA-binding domain, leads to loss of Zld occupancy at CAGGTA sites, localization of Zld to new loci, and the formation of stabilized protein hubs (Fallacaro et al. 2024). The authors suggest that that these changes in Zld occupancy, and that of other transcription factors, are dictated by both the DNA-binding activity and by interactions of non-DNA-binding domains with cofactors, like GAF. Counter to the model that binding is driven by interactions between unstructured domains and cofactors, our data suggest that Zld binding is driven by differential strengths of two distinct DNA-binding domains. In culture, GAF is not required for Zld occupancy as knockdown of GAF does not lead to a change in the binding of either wild-type Zld or Zld^ZnF2^. Our analysis of the embryonic binding sites to which Zld localizes upon mutation of ZnF5 shows that these are regions bound by Zld in neuroblasts, suggesting that when the CAGGTA-binding domain is disrupted ZnF2 results in the observed redistribution of Zld binding. We suggest that future studies should consider that even largely disordered proteins may harbor small, structured domains with unanticipated functional roles.

Tissue-specific regulation of pioneer activity may be governed by how a protein interacts with chromatin. Even though Zld^ZnF3-6^ binds chromatin via ZnF2, this occupancy is not able to promote chromatin accessibility. This demonstrates that the CAGGTA-binding domain is required for pioneer activity separate from chromatin binding. Furthermore, Zld bound regions in neuroblasts enriched for the CAGGTA motif lose accessibility promptly upon exit from the multipotent state, coincident with lack of Zld. By contrast, Zld-bound regions enriched for the G-rich motif lose accessibility gradually as cells differentiate, suggesting that other factors maintain accessibility in the absence of Zld. We propose that at these regions, Zld may not function as a pioneer factor but instead may promote transcription through mechanisms more similar to other transcriptional activators.

We propose a model in which Zld function is regulated by two DNA-binding domains, with domain choice dictated by developmental context. We suggest that this mechanism may not be unique to Zld. Indeed POU transcription factors, like Oct4, contain both a POU-specific domain and a POU-homeodomain connected by a flexible linker (Malik et al. 2018; Carminati et al. 2024). The discovery of a previously unrecognized DNA-binding domain in Zld, despite over a decade of study, underscores the importance of examining pioneer-factor function across developmental contexts. Future insights into the mechanisms governing the tissue-specific usage of distinct binding domains will also impact our understanding of how chimeric oncoproteins that juxtapose distinct DNA-binding or effector domains create new combinations with emergent and sometimes antagonistic regulatory properties. Together, our data suggest that the modularity provided by multiple DNA-binding domains may allow a flexible mechanism to achieve tissue-specific outcomes and indicate broader principles governing pioneer-factor biology.

## Materials and Methods

### Drosophila strains

Flies were raised on standard fly food at 25°C (unless otherwise noted). HA-tagged constructs were cloned into pUAST-attB (Addgene). Transgenes were inserted into the *pUAST-attB M{3xP3- RFP.attP}ZH-86Fb* docking site using ΦC31 integrase-mediated transgenesis (BestGene, Chino Hills, CA). For ectopic expression of Zld in brains, HA-tagged versions of Zld with mutations in ZnF2, ZnF3-6 or ZnF2-6, were collected from *brat^11/DF,Tub-Gal80ts^*^;*UAS-HA-transgene*^ larvae, in which *Worniu-Gal4* drives overexpression of HA-Zld transgenes in neuroblasts. Larvae were grown at 18°C until they reached L3 stage and then they were shifted to 37°C for 24 hours to induce ectopic expression of the HA-Zld constructs. GFP-negative and non-tubby larvae were selected for brains dissections.

### Cell culture, generation of stable cell lines, luciferase assays and RNAi treatment

The stable cell line expressing Zld WT used in this study was previously described (Gibson et al. 2024). *zld* cDNA containing mutations in ZnF2, ZnF3-6 or ZnF2-6 was cloned into pMT-puro (Addgene) by Gibson assembly (New England Biolabs) as previously described (Gibson et al. 2024) and cells were continuously selected with 1 µg/ml of puromycin. Zld expression was induced by adding 1000uM CuSO_4_ to the cell culture medium. For luciferases assays, plasmids expressing *zld* cDNA with mutations in ZnF2 and ZnF3-6 under the *actin* promoter (pAc5.1) (Hamm et al. 2017, 2015) were transiently transfected in triplicate into 24-well dishes with a total of 300 ng of DNA using Effectene transfection reagent (Qiagen), as previously described. Luciferase assays were performed using the Dual-Luciferase assay system (Promega). Fold activation was determined by comparison with transfections using pAc5.1 alone.

We generated dsRNA for GAF and LacZ by PCR amplifying a dsDNA with T7 RNA polymerase promoters on the 5’ end of both strands and then generated dsRNA using T7 RNA polymerase (New England BioLabs). Oligonucleotide primer sequences were obtained from Judd et al., 2020. All RNAi treatments were done as previously described (Judd et al. 2020). Briefly, cells were treated with 10 µg/mL for three days, then split into two wells. Fresh media and additional dsRNA were added to maintain a final concentration of 10 µg/mL, and incubation was continued for two additional days. After the initial three days of RNAi treatment, cells were induced with 1000 µM CuSO₄ for 48 hours and subsequently harvested.

### Immunoblotting

Proteins were transferred to 0.45 μm Immobilon-P PVDF membrane (Millipore, Burlington, MA) in transfer buffer (25 mM Tris, 200 mM Glycine, 20% methanol) for 75 min at 500mA at 4°C. Membranes were blocked with BLOTTO (2.5% non-fat dry milk, 0.5% BSA, 0.5% NP-40, in TBST) for 30 min at room temperature and then incubated with anti-Zld (1:750) (Harrison et al. 2010), anti-GAF (1:100) (Gaskill et al. 2023), anti-HA–peroxidase (1:500) (clone 3F10, Roche), anti-tubulin (1:5,000) (DM1A, Sigma), V5 (14440-1-AP, Proteintech) overnight at 4 °C. The secondary incubation was performed with goat anti-rabbit IgG-HRP conjugate (1:3000) (Bio-Rad) or anti-mouse IgG-HRP conjugate (1:3000) (Bio-Rad) for 1 hr at room temperature. Blots were treated with SuperSignal West Pico PLUS chemiluminescent substrate (Thermo Fisher Scientific) and visualized using the Azure Biosystems c600.

### Protein Purification

Zld N (Zld^1-619^), Zld N (Zld^1-619(ZnF2)^) and Zld C (Zld^1117–1487^) were cloned into pGEX-6P-1, a T7-driven GST expression vector (Addgene). GST-Zld N, GST-Zld N^ZnF2^ and Zld C were purified from *E. coli* in column buffer (20mM Hepes (pH 7.6), 0.5M KCl, 1mM EDTA, 10mM DTT and 0.8mM PMSF and 0.01% NP40) using GST beads (New England Biolabs). After washing, protein was eluted with elution buffer (20mM Hepes (pH 7.6), 0.150 M KCl, 1 mM EDTA, 10 mM DTT and 0.8 mM PMSF and 0.2% NP40.

### Electromobility Shift Assays

EMSAs were performed as previously described (McDaniel et al. 2019). Cy5-labelled probes was incubated with recombinant proteins (Zld N, Zld N^ZnF2^ and Zld C) in buffer containing: 25ng poly[d-(IC)], 12.5mM HEPES, 0.5mM EDTA, 0.5mM EGTA, 5% glycerol, 93.75μM ZnCl2, 0.375mM DTT, 75μM PMSF, 0.075mg/ml BSA, 5mM MgCl2, 0.00625% NP-40 and 50mM KCl at room temperature for 60 min. 12.5 and 25nM of recombinant protein (Zld N or Zld C) were used in each reaction with three different probes and 3, 9 and 12.5nM recombinant protein (Zld N or Zld N^ZnF2^) was used in each reaction with the G-rich probe. The sequence of the oligonucleotides probes that were Cy5 labeled were as follows: GAGAGAGA<u>CTACCTG</u>TGGCTCACT (wild type), GAGAGAGA<u>GTAGTTC</u>TGGCTCACT (mutant) and GAGAGAGA<u>TTGGGAA</u>TGGCTCACT (AAGGGT).

### Protein Binding Microarrays

For PBMs, GST-tagged constructs expressing either Zld N^103-608^ and Zld C^1241-1596^ proteins were used as previously described (Brennan et al. 2023). Briefly, proteins were expressed using the PURExpress In Vitro Protein Synthesis Kit (New England BioLabs) and assayed in duplicate on two PBM arrays containing distinct probe sequences. Brujin sequences were used to construct universal arrays to provide an unbiased representation of all possible 7-mers DNA sequences. Fluorescence intensity data were processed to extract binding signals and used to generate DNA-binding motifs with Top10AlignZ (Weirauch et al. 2014). For quantitative analysis, E-scores and Z-scores were calculated for every 7-mer as previously (Weirauch et al. 2014; Berger et al. 2006).

### Cleavage Under Targets & Release Using Nuclease (CUT&RUN)

Fifty GFP- and non tubby larval brains were dissected into Schneider’s medium, transfer and dounced into Wash Buffer (20 mM HEPES, 150mM NaCl and 0.5mM spermidine) supplemental with 0.1% BSA as described in CUTANA CUT&RUN (Epicypher) ten times using a tight pestle. The Epicypher protocol was followed with the following modifications: the Digitonin Buffer was prepared at 0.05% concentration; DNA cleanup was performed using the MinElute Reaction Cleanup Kit (Qiagen); and libraries were prepared using the NEBNext Ultra II DNA Library Prep Kit (E7645S, New England BioLabs), with adaptors diluted 1:30. AMPure XP beads (Beckman Coulter) were used at 1.1X ratio following adaptor ligation and 0.9X following PCR. PCR cycling parameters followed those outlined in the Epicypher protocol. Antibodies used included 1µL of anti-Zld and 1µL of anti-HA (12CA5; Sigma-Aldrich). Libraries were sequenced on an Illumina NovaSeq 6000 platform using 150 bp paired-end reads at the UW-Madison Biotechnology Center.

### CUT&RUN Analysis

Adapters and low-quality bases were removed using Trimmomatic-0.39 (Bolger et al. 2014). Reads were aligned to the *Drosophila melanogaster* reference genome (version dm6) using bowtie 2 v2.3.5 (Lawrence et al. 2013). Unmapped, multiply aligning, mitochondrial, and scaffold reads were removed. To identify regions that were enriched in immunoprecipitated samples relative to input controls, peak calling was performed using MACS v2 (Zhang et al. 2008) with the following parameters: -g 1.2e8, --call-summits. From experiments using Zld antibody peaks were called with respect to a rab-IgG CUT&RUN control. From experiments using HA antibody peaks were called with respect to a HA CUT&RUN experiment in a *brat* mutant brain (no transgene overexpression). To focus analysis on robust, high-quality peaks, we used 100 bp up- and downstream of peak summits, and regions belonging to contigs and unmapped chromosomes were removed. z score-normalized bigWigs were created by subtracting the mean read coverage (counts) from merged replicate read counts in 10bp bins across the entire genome and dividing by the standard deviation. Heatmaps and average signal line plots were generated using deepTools2 (Ramírez et al. 2016) with z score-normalized bigWig files.

### ChIP-seq

ChIP-seq experiments were performed on hand-sorted stage 5 embryos laid by *Zld*^ZnF2^ homozygous females, as previously described (Hamm et al. 2017), as well as on *w^1118^* embryos as a wild-type control.

For each genotype, two biological replicates were collected, with one thousand embryos per replicate. Briefly, embryos were dechorionated in 50% bleach for 3 minutes and fixed in 0.45% formaldehyde. They were then homogenized in RIPA buffer (50 mM Tris-Cl pH 8.0, 0.1% SDS, 1% Triton-100, 0.5% deoxycholate and 150 mM NaCl), and all subsequent steps were performed as previously described (Marsh et al. 2024, Gaskill et al 2022). Spike-in (5%) chromatin of fixed and sonicated *Drosophila virilis* was added for immunoprecipitation. For each ChIP-seq experiment in S2 cells, ∼25 million cells were cross-linked and processed following the protocol previously described (Gibson et al. 2024). For each genotype, two biological replicates were collected. Preparation of sequencing libraries of both Chip-seq experiments in embryos and S2 cells was performed using the NEBNext Ultra II kit (NEB) with eight PCR cycles for library amplification. Sequencing was performed on an Illumina Novaseq 6000 instrument using 50-bp single-end reads.

### ChIP-seq Analysis

Adapters and low-quality bases were removed using Trimmomatic-0.39 (Bolger et al. 2014). Reads were aligned to the *Drosophila melanogaster* reference genome (version dm6) using bowtie 2 v2.3.5 (Lawrence et al. 2013). Unmapped, multiply aligning, mitochondrial, and scaffold reads were removed. To identify regions that were enriched in immunoprecipitated samples relative to input controls, peak calling was performed using MACS v2 (Zhang et al. 2008) with the following parameters: -g 1.2e8, --call-summits. The BEDtools intersectBed function was used to define high confidence peaks present in both replicates(Quinlan and Hall 2010). To focus analysis on robust, high-quality peaks, we used 100 bp up- and downstream of peak summits, and regions belonging to contigs and unmapped chromosomes were removed. All bigwigs were generated using bamCoverage from deepTools v3.5.192. For ChIP-seq experiment in S2 cells z-score normalized Bigwigs were created by taking the average read count across all bins genome wide and dividing by the standard deviation. IPs for Zld in the embryo were spike-in normalized in which a scaling factor was determined by the ratio of percent *D. virilis* chromatin aligned in the IPs as compared to input. Heatmaps and average signal line plots were generated using deepTools2 (Ramírez et al. 2016) with z score-normalized (for S2 cells) or spike-in normalized (for embryos) bigWig files.

### ATAC-seq

For each ATAC-seq experiment in S2 cells, ∼200 thousand cells were processed following the protocol previously described Gibson et al. 2024. For each genotype, two biological replicates were collected. Samples were tagmented with 2.5 µl of Tagment DNA Enzyme (Illumina) and incubated at 37 °C for 30 min. After DNA tagmentation, DNA was purified using the MinElute Reaction Cleanup kit (Qiagen) and eluted in 10 µl of buffer EB. DNA was PCR amplified for twelve cycles. Amplified libraries were purified using a 1.2× ratio of Axygen paramagnetic beads. Sequencing was performed on an Illumina NovaSeq X Plus instrument using 150-bp pair-end reads.

### ATAC-seq analysis

Adaptor sequences were removed from raw ATAC-seq reads using NGmerge (v0.3). The resulting reads were aligned to the *D. melanogaster* genome (dm6) with Bowtie2 using -k 2 --very-sensitive --no-mixed --no-discordant -X 5000. Reads with a mapping quality below 30, as well as those mapping to the mitochondrial genome or scaffolds, were excluded from further analysis. As described previously, only fragments <100 bp in length were considered for downstream analysis (Gibson 2024). Reads from all replicates were merged, and MACS2 was employed for peak calling with -f BAMPE --keep-dup all -g dm --call-summits. featureCounts was used to quantify reads within 200 bp of peak summits (±100 bp). Differential chromatin accessibility was determined using DESeq2, and regions with an adjusted *P* value <0.05 were considered significantly differentially accessible (Love et al. 2014).

### Immunostaining

Third instar larval brains were dissected in solution containing 4 % formaldehyde, 1xPBS and fixed in 100 mM PIPES (pH6.9), 1 mM EGTA, 0.3 % Triton X-100, and 1 mM MgSO4 for 23 min at room temperature. Fixed brains were washed with 0.3 % Triton X-100 in 1x PBS (PBST). After removing fix solution, samples were incubated with primary antibodies for 3 hours at room temperature. Samples were washed with PBST and then incubated with secondary antibodies and Phalloidin (ThermoFisher Scientific) overnight at 4°C. On the next day, samples were washed with PBST and then equilibrated in Prolong Gold antifade mount (ThermoFisher Scientific). Brains were mounted with dorsal surface up. Confocal images were acquired on a Leica SP5 scanning confocal microscope (Leica Microsystems Inc). To quantify the number of type II neuroblasts, images of brain lobes were sequentially taken at every 1.5 mm from dorsal to ventral. Ten brains per genotype were used to obtain data.

## Acknowledgements

Stocks obtained from the Bloomington *Drosophila* Stock Center (National Institutes of Health P40OD018537) were used in this study. Flybase (FB2024_02) was used throughout the study. This work was supported by National Institutes of Neurological Disease and Stroke grants R01NS111647(to MMH and CYL); National Institutes of General Medical Sciences grant R35GM136298 (to MMH), R35GM148241(to CAR). ETZ was supported by a Postdoctoral Fellowship from the University of Wisconsin-Madison Stem Cell and Regenerative Medicine Center.

## Author Contributions

ETZ, MMH, HK, CYL and CAR conceived the study. Most experiments were designed and analyzed by ETZ, HK, and MMH. H-YL, CAR, AY, and TRH performed protein-binding array analysis. EDL and ZAF generated and analyzed transgenic fly lines. ETZ and MMH wrote the original draft of the manuscript. HK, CYL, and CAR reviewed and edited the manuscript. MMH, CYL, CAR and TRH acquired funding.

## Supplemental Figures

**Supplementary Fig. 1: GAF does not drive Zld redistribution in S2 cells when ZnF2 is mutated. a.** Percentage of Zld ChIP-seq peaks unique to wild-type Zld unique, unique to Zld^ZnF3-6^ or shared that contain the GAGA motif. **b.** Percentage of ChIP-seq peaks unique to wild-type Zld unique, unique to Zld^ZnF2^ or shared that contain the GAGA motif **c.** Immunoblot of Zld or GAF as indicated from S2 cells induced for Zld expression and treated with either LacZ or GAF dsRNA. Tubulin is shown as a loading control. **d.** Metaplots of Zld ChIP-seq from S2 cells induced for Zld expression and treated with either LacZ or GAF dsRNA. Signal is centered on Zelda peaks. **e-f.** Heat maps of z-score normalized Zld ChIP-seq centered on peak summits with 500 bp flanking sequence and ordered by signal in wild-type Zld. Genotypes are indicated above, and peak classes are indicated on the left.

**Supplementary Fig. 2: HA-tagged Zld constructs recapitulate Zld genomic binding and activity than untagged-Zld versions. a.** Immunostaining of third instar larval brains lobes expressing ectopic Zld or HA-Zld constructs as indicated. Expression is driven in type I and type II neuroblasts by *Wor-Gal4*, *Tub-Gal80^ts^*. Yellow arrows indicate a selection of Dpn+, Ase- type II neuroblasts. Scale bar, 20µm. **b.** Number of type II neuroblasts per lobe of brains in which the indicated Zld and HA-Zld constructs are expressed. For each genotype: n = 10 brains. ****, p < 0.0001, ***, p < 0.001, **, p < 0.01. ns= not significant as determined by ANOVA. **c.** Immunoblots for HA-tagged proteins in brains dissected from the genotypes indciated. Tubulin is shown as a loading control. **d.** Venn diagram of peaks from anti-Zld CUT&RUN from *brat^11/Df^* brains and *brat^11/Df^* brains expressing HA-Zld. **e.** Venn diagram of peaks from anti-Zld CUT&RUN from *brat^11/Df^* brains and anti-HA CUT&RUN from *brat^11/Df^* brains expressing HA-Zld.

**Supplementary Fig.3: ZnF2 is dispensable for Zld binding in the embryo, but mediates redistribution to regions bound by Zld in neuroblasts. a.** Immunoblot for Zld on stage 5 embryos demonstrate equivalent protein levels for wild type and ZnF2 mutant. Tubulin is shown as a loading control. **b.** Heatmaps of Zld ChIP-seq in wild-type and Zld^ZnF2^ embryos centered on Zld-bound peaks. **c.** Genome browser tracks showing representative loci of up-regulated Zld- target genes from Hamm et al. *PLoS Genet* 2017. Zld binding in wild-type and Zld^ZnF2^ embryos along with CAGGTA motifs are shown. **d.** Heatmaps of Zld occupancy in embryos and neuroblasts for the genotypes indicated above at genomic sites defined by comparing wild-type Zld and Zld^ZnF5^ in the embryo (from Fallacaro et al. 2024).

**Supplementary Fig.4: The N-terminus of Zld is sufficient for mediating transcriptional activation. a.** Cartoon schematic of VP16-Zld constructs. **b.** Immunostaining of third instar larval brains lobes expressing ectopic Zld or VP16-NLS constructs as indicated. Expression is driven in type I and type II neuroblasts by *Wor-Gal4*, *Tub-Gal80^ts^*. Yellow arrows indicate a selection of Dpn+, Ase- type II neuroblasts. Scale bar, 20µm. **c.** Luciferase reporter assays with full-length Zld, Zld^1-619^ WT and ZnF2 mutant (Zld1-619^(ZnF2)^). Data are shown for both the scute reporter with the four CAGGTA replaced with AAGGGT and the neuroblast-specific *tll* enhancer. Immunoblots for V5 tagged proteins indicate protein expression levels. Tubulin is shown as a loading control.

## References

Andersson R, Sandelin A. 2020. Determinants of enhancer and promoter activities of regulatory elements. Nat Rev Genet 21: 71–87.

Balsalobre A, Drouin J. 2022. Pioneer factors as master regulators of the epigenome and cell fate. Nat Rev Mol Cell Biol 23: 449–464.

Barral A, Zaret KS. 2024. Pioneer factors: roles and their regulation in development. Trends in Genetics 40: 134–148.

Berger MF, Philippakis AA, Qureshi AM, He FS, Estep PW, Bulyk ML. 2006. Compact, universal DNA microarrays to comprehensively determine transcription-factor binding site specificities. Nat Biotechnol 24: 1429–1435.

Bolger AM, Lohse M, Usadel B. 2014. Trimmomatic: a flexible trimmer for Illumina sequence data. Bioinformatics 30: 2114–2120.

Bosch JRT, Benavides JA, Cline TW. 2006. The TAGteam DNA motif controls the timing of *Drosophila* pre-blastoderm transcription. Development 133: 1967–1977.

Brennan KJ, Weilert M, Krueger S, Pampari A, Liu H, Yang AWH, Morrison JA, Hughes TR, Rushlow CA, Kundaje A, et al. 2023. Chromatin accessibility in the Drosophila embryo is determined by transcription factor pioneering and enhancer activation. Developmental Cell 58: 1898–1916.e9.

Buecker C, Srinivasan R, Wu Z, Calo E, Acampora D, Faial T, Simeone A, Tan M, Swigut T, Wysocka J. 2014. Reorganization of Enhancer Patterns in Transition from Naive to Primed Pluripotency. Cell Stem Cell 14: 838–853.

Carminati M, Vecchia L, Stoos L, Thomä NH. 2024. Pioneer factors: Emerging rules of engagement for transcription factors on chromatinized DNA. Current Opinion in Structural Biology 88: 102875.

Chronis C, Fiziev P, Papp B, Butz S, Bonora G, Sabri S, Ernst J, Plath K. 2017. Cooperative Binding of Transcription Factors Orchestrates Reprogramming. Cell 168: 442–459.e20.

Donaghey J, Thakurela S, Charlton J, Chen JS, Smith ZD, Gu H, Pop R, Clement K, Stamenova EK, Karnik R, et al. 2018. Genetic determinants and epigenetic effects of pioneer-factor occupancy. Nat Genet 50: 250–258.

Fallacaro S, Mukherjee A, Ratchasanmuang P, Zinski J, Haloush YI, Shankta K, Mir M. 2024. A fine kinetic balance of interactions directs transcription factor hubs to genes. bioRxiv 2024.04.16.589811.

Ferrie JJ, Karr JP, Tjian R, Darzacq X. 2022. “Structure”-function relationships in eukaryotic transcription factors: The role of intrinsically disordered regions in gene regulation. Molecular Cell 82: 3970– 3984.

Foo SM, Sun Y, Lim B, Ziukaite R, O’Brien K, Nien C-Y, Kirov N, Shvartsman SY, Rushlow CA. 2014. Zelda Potentiates Morphogen Activity by Increasing Chromatin Accessibility. Current Biology 24: 1341–1346.

Freund MM, Harrison MM, Torres-Zelada EF. 2024. Exploring the reciprocity between pioneer factors and development. Development 151: dev201921.

Gaskill MM, Gibson TJ, Larson ED, Harrison MM. 2021. GAF is essential for zygotic genome activation and chromatin accessibility in the early Drosophila embryo. eLife 10: e66668.

Gaskill MM, Soluri IV, Branks AE, Boka AP, Stadler MR, Vietor K, Huang H-YS, Gibson TJ, Mukherjee A, Mir M, et al. 2023. Localization of the Drosophila pioneer factor GAF to subnuclear foci is driven by DNA binding and required to silence satellite repeat expression. Developmental Cell 58: 1610–1624.e8.

Gibson TJ, Larson ED, Harrison MM. 2024. Protein-intrinsic properties and context-dependent effects regulate pioneer factor binding and function. Nat Struct Mol Biol 31: 548–558.

Hamm DC, Bondra ER, Harrison MM. 2015. Transcriptional Activation Is a Conserved Feature of the Early Embryonic Factor Zelda That Requires a Cluster of Four Zinc Fingers for DNA Binding and a Low-complexity Activation Domain. Journal of Biological Chemistry 290: 3508–3518.

Hamm DC, Larson ED, Nevil M, Marshall KE, Bondra ER, Harrison MM. 2017. A conserved maternal-specific repressive domain in Zelda revealed by Cas9-mediated mutagenesis in Drosophila melanogaster ed. M.B. Eisen. PLoS Genet 13: e1007120.

Harrison MM, Botchan MR, Cline TW. 2010. Grainyhead and Zelda compete for binding to the promoters of the earliest-expressed Drosophila genes. Developmental biology 345: 248–255.

Harrison MM, Li X-YY, Kaplan T, Botchan MR, Eisen MB. 2011. Zelda binding in the early Drosophila melanogaster embryo marks regions subsequently activated at the maternal-to-zygotic transition ed. G.P. Copenhaver. PLoS Genet 7: e1002266.

Heinz S, Romanoski CE, Benner C, Glass CK. 2015. The selection and function of cell type-specific enhancers. Nat Rev Mol Cell Biol 16: 144–154.

Homem CCF, Knoblich JA. 2012. Drosophila neuroblasts: a model for stem cell biology. Development 139: 4297–4310.

Horisawa K, Suzuki A. 2023. The role of pioneer transcription factors in the induction of direct cellular reprogramming. Regenerative Therapy 24: 112–116.

Iwafuchi-Doi M, Zaret KS. 2014. Pioneer transcription factors in cell reprogramming. Genes Dev 28: 2679–2692.

Janssens DH, Komori H, Grbac D, Chen K, Koe CT, Wang H, Lee C-Y. 2014. Earmuff restricts progenitor cell potential by attenuating the competence to respond to self-renewal factors. Development 141: 1036–1046.

Kobayashi W, Tachibana K. 2021. Awakening of the zygotic genome by pioneer transcription factors. Current Opinion in Structural Biology 71: 94–100.

Kumar DK, Jonas F, Jana T, Brodsky S, Carmi M, Barkai N. 2023. Complementary strategies for directing in vivo transcription factor binding through DNA binding domains and intrinsically disordered regions. Molecular Cell 83: 1462–1473.e5.

Larson ED, Komori H, Gibson TJ, Ostgaard CM, Hamm DC, Schnell JM, Lee C-Y, Harrison MM. 2021a. Cell-type-specific chromatin occupancy by the pioneer factor Zelda drives key developmental transitions in Drosophila. Nat Commun 12: 7153.

Larson ED, Marsh AJ, Harrison MM. 2021b. Pioneering the developmental frontier. Molecular Cell 81: 1640–1650.

Lawrence M, Huber W, Pagès H, Aboyoun P, Carlson M, Gentleman R, Morgan MT, Carey VJ. 2013. Software for Computing and Annotating Genomic Ranges ed. A. Prlic. PLoS Comput Biol 9: e1003118.

Lee CS, Friedman JR, Fulmer JT, Kaestner KH. 2005. The initiation of liver development is dependent on Foxa transcription factors. Nature 435: 944–947.

Lee C-Y, Wilkinson BD, Siegrist SE, Wharton RP, Doe CQ. 2006. Brat Is a Miranda Cargo Protein that Promotes Neuronal Differentiation and Inhibits Neuroblast Self-Renewal. Developmental Cell 10: 441–449.

Lee K, Cho H, Rickert RW, Li QV, Pulecio J, Leslie CS, Huangfu D. 2019. FOXA2 Is Required for Enhancer Priming during Pancreatic Differentiation. Cell Reports 28: 382–393.e7.

Li X, MacArthur S, Bourgon R, Nix D, Pollard DA, Iyer VN, Hechmer A, Simirenko L, Stapleton M, Hendriks CLL, et al. 2008. Transcription Factors Bind Thousands of Active and Inactive Regions in the Drosophila Blastoderm ed. J. Kadonaga. PLoS Biol 6: e27.

Liang HL, Nien CY, Liu HY, Metzstein MM, Kirov N, Rushlow C. 2008. The zinc-finger protein Zelda is a key activator of the early zygotic genome in Drosophila. Nature 456: 400–403.

Love MI, Huber W, Anders S. 2014. Moderated estimation of fold change and dispersion for RNA-seq data with DESeq2. Genome Biol 15: 550.

Malik V, Zimmer D, Jauch R. 2018. Diversity among POU transcription factors in chromatin recognition and cell fate reprogramming. Cell Mol Life Sci 75: 1587–1612.

Mayran A, Khetchoumian K, Hariri F, Pastinen T, Gauthier Y, Balsalobre A, Drouin J. 2018. Pioneer factor Pax7 deploys a stable enhancer repertoire for specification of cell fate. Nat Genet 50: 259–269.

McDaniel SL, Gibson TJ, Schulz KN, Fernandez Garcia M, Nevil M, Jain SU, Lewis PW, Zaret KS, Harrison MM. 2019. Continued Activity of the Pioneer Factor Zelda Is Required to Drive Zygotic Genome Activation. Molecular Cell 74: 185–195.e4.

Nien C-Y, Liang H-L, Butcher S, Sun Y, Fu S, Gocha T, Kirov N, Manak JR, Rushlow C. 2011. Temporal Coordination of Gene Networks by Zelda in the Early Drosophila Embryo ed. G.S. Barsh. PLoS Genet 7: e1002339.

Quinlan AR, Hall IM. 2010. BEDTools: a flexible suite of utilities for comparing genomic features. Bioinformatics 26: 841–842.

Rada-Iglesias A, Bajpai R, Swigut T, Brugmann SA, Flynn RA, Wysocka J. 2011. A unique chromatin signature uncovers early developmental enhancers in humans. Nature 470: 279–283.

Rajan A, Ostgaard CM, Lee C-Y. 2021. Regulation of Neural Stem Cell Competency and Commitment during Indirect Neurogenesis. Int J Mol Sci 22: 12871.

Ramírez F, Ryan DP, Grüning B, Bhardwaj V, Kilpert F, Richter AS, Heyne S, Dündar F, Manke T. 2016. deepTools2: a next generation web server for deep-sequencing data analysis. Nucleic acids research 44: W160–W165.

Reichardt I, Bonnay F, Steinmann V, Loedige I, Burkard TR, Meister G, Knoblich JA. 2018. The tumor suppressor Brat controls neuronal stem cell lineages by inhibiting Deadpan and Zelda. EMBO Reports 19: 102–117.

Ribeiro L, Tobias-Santos V, Santos D, Antunes F, Feltran G, de Souza Menezes J, Aravind L, Venancio TM, Nunes da Fonseca R. 2017. Evolution and multiple roles of the Pancrustacea specific transcription factor zelda in insects ed. C. Desplan. PLOS Genetics 13: e1006868.

Rives-Quinto N, Komori H, Ostgaard CM, Janssens DH, Kondo S, Dai Q, Moore AW, Lee C-Y. 2020. Sequential activation of transcriptional repressors promotes progenitor commitment by silencing stem cell identity genes. eLife 9: e56187.

Schulz KN, Bondra ER, Moshe A, Villalta JE, Lieb JD, Kaplan T, McKay DJ, Harrison MM. 2015. Zelda is differentially required for chromatin accessibility, transcription factor binding, and gene expression in the early *Drosophila* embryo. Genome Research 25: 1715–1726.

Schulz KN, Harrison MM. 2019. Mechanisms regulating zygotic genome activation. Nature Reviews Genetics 20: 221–234.

Shen Y, Liu K, Liu J, Shen J, Ye T, Zhao R, Zhang R, Song Y. 2025. TBP bookmarks and preserves neural stem cell fate memory by orchestrating local chromatin architecture. Molecular Cell 85: 413–429.e10.

Soufi A, Donahue G, Zaret KS. 2012. Facilitators and Impediments of the Pluripotency Reprogramming Factors’ Initial Engagement with the Genome. Cell 151: 994–1004.

Sun Y, Nien C-Y, Chen K, Liu H-Y, Johnston J, Zeitlinger J, Rushlow C. 2015. Zelda overcomes the high intrinsic nucleosome barrier at enhancers during Drosophila zygotic genome activation. Genome research 25: 1703–14.

Vastenhouw NL, Cao WX, Lipshitz HD. 2019. The maternal-to-zygotic transition revisited. Development 146: dev161471.

Veil M, Yampolsky LY, Grüning B, Onichtchouk D. 2019. Pou5f3, SoxB1, and Nanog remodel chromatin on high nucleosome affinity regions at zygotic genome activation. Genome Res 29: 383–395.

Weirauch MT, Yang A, Albu M, Cote AG, Montenegro-Montero A, Drewe P, Najafabadi HS, Lambert SA, Mann I, Cook K, et al. 2014. Determination and Inference of Eukaryotic Transcription Factor Sequence Specificity. Cell 158: 1431–1443.

Xu Z, Chen H, Ling J, Yu D, Struffi P, Small S. 2014. Impacts of the ubiquitous factor Zelda on Bicoid-dependent DNA binding and transcription in *Drosophila*. Genes Dev 28: 608–621.

Yáñez-Cuna JO, Dinh HQ, Kvon EZ, Shlyueva D, Stark A. 2012. Uncovering *cis* -regulatory sequence requirements for context-specific transcription factor binding. Genome Res 22: 2018–2030.

Yartseva V, Giraldez AJ. 2015. The Maternal-to-Zygotic Transition During Vertebrate Development: A Model for Reprogramming. Curr Top Dev Biol 113: 191–232.

Zaret KS. 2020. Pioneer Transcription Factors Initiating Gene Network Changes. Annu Rev Genet 54: 367–385.

Zhang Y, Liu T, Meyer CA, Eeckhoute J, Johnson DS, Bernstein BE, Nusbaum C, Myers RM, Brown M, Li W, et al. 2008. Model-based Analysis of ChIP-Seq (MACS). Genome Biol 9: R137.

