## Supplemental Figures for "Differential usage of two, distinct DNA-binding domains regulates tissue-specific occupancy of the pioneer factor Zelda"

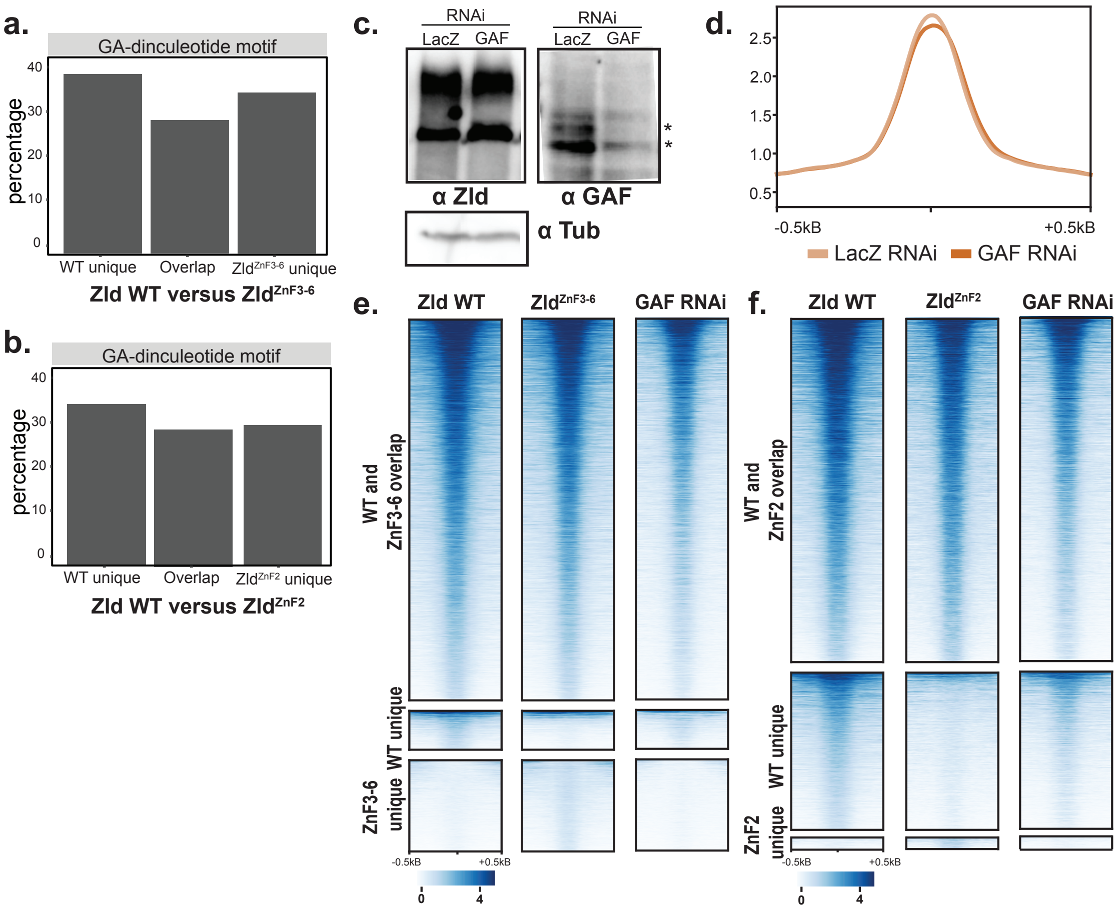


**Supplementary Fig. 1: GAF does not drive Zld redistribution in S2 cells when ZnF2 is mutated**. **a.** Percentage of Zld ChIP-seq peaks unique to wild-type Zld unique, unique to Zld$^{ZnF3-6}$ or shared that contain the GAGA motif. b**.** Percentage of ChIP-seq peaks unique to wild-type Zld unique, unique to Zld$^{ZnF2}$ or shared that contain the GAGA motif **c.** Immunoblot of Zld or GAF as indicated from S2 cells induced for Zld expression and treated with either LacZ or GAF dsRNA. Tubulin is shown as a loading control. **d.** Metaplots of Zld ChIP-seq from S2 cells induced for Zld expression and treated with either LacZ or GAF dsRNA. Signal is centered on Zelda peaks. **e-f.** Heat maps of z-score normalized Zld ChIP-seq centered on peak summits with 500 bp flanking sequence and ordered by signal in wild-type Zld. Genotypes are indicated above, and peak classes are indicated on the left.

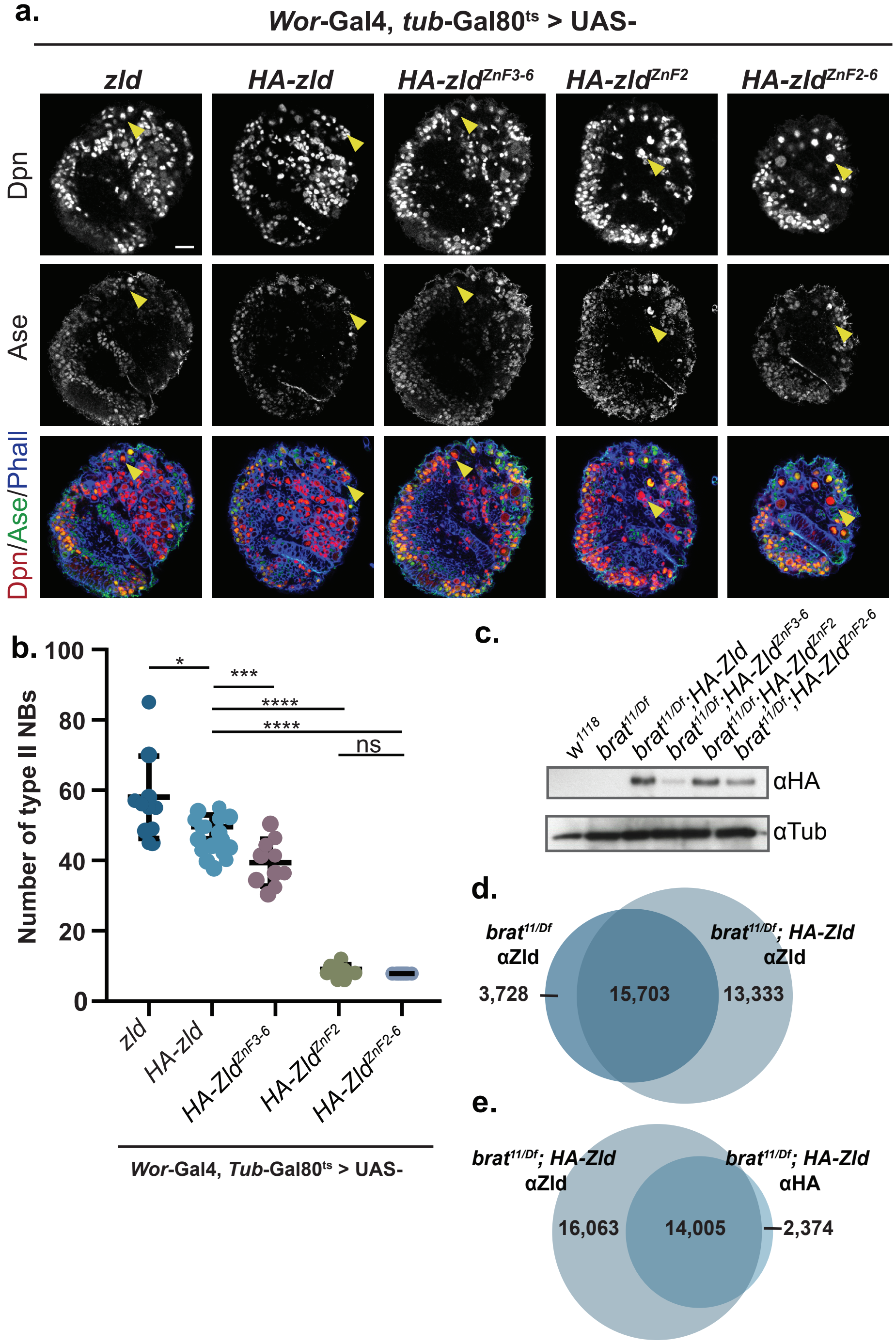


**Supplementary Fig. 2: HA-tagged Zld constructs recapitulate Zld genomic binding and activity than untagged-Zld versions. a.** Immunostaining of third instar larval brains lobes expressing ectopic Zld or HA-Zld constructs as indicated. Expression is driven in type I and type II neuroblasts by *Wor-Gal4, Tub-Gal80*$^{ts}$. Yellow arrows indicate a selection of Dpn+, Ase- type II neuroblasts. Scale bar, 20µm. **b.** Number of type II neuroblasts per lobe of brains in which the indicated Zld and HA-Zld constructs are expressed. For each genotype: n = 10 brains. ****, $p < 0.0001$, ***, $p < 0.001$, **, $p < 0.01$. ns= not significant as determined by ANOVA. **c.** Immunoblots for HA-tagged proteins in brains dissected from the genotypes indciated. Tubulin is shown as a loading control. **d.** Venn diagram of peaks from anti-Zld CUT&RUN from *brat*$^{11/Df}$ brains and *brat*$^{11/Df}$ brains expressing HA-Zld. **e.** Venn diagram of peaks from anti-Zld CUT&RUN from *brat*$^{11/Df}$ brains and anti-HA CUT&RUN from *brat*$^{11/Df}$ brains expressing HA-Zld.

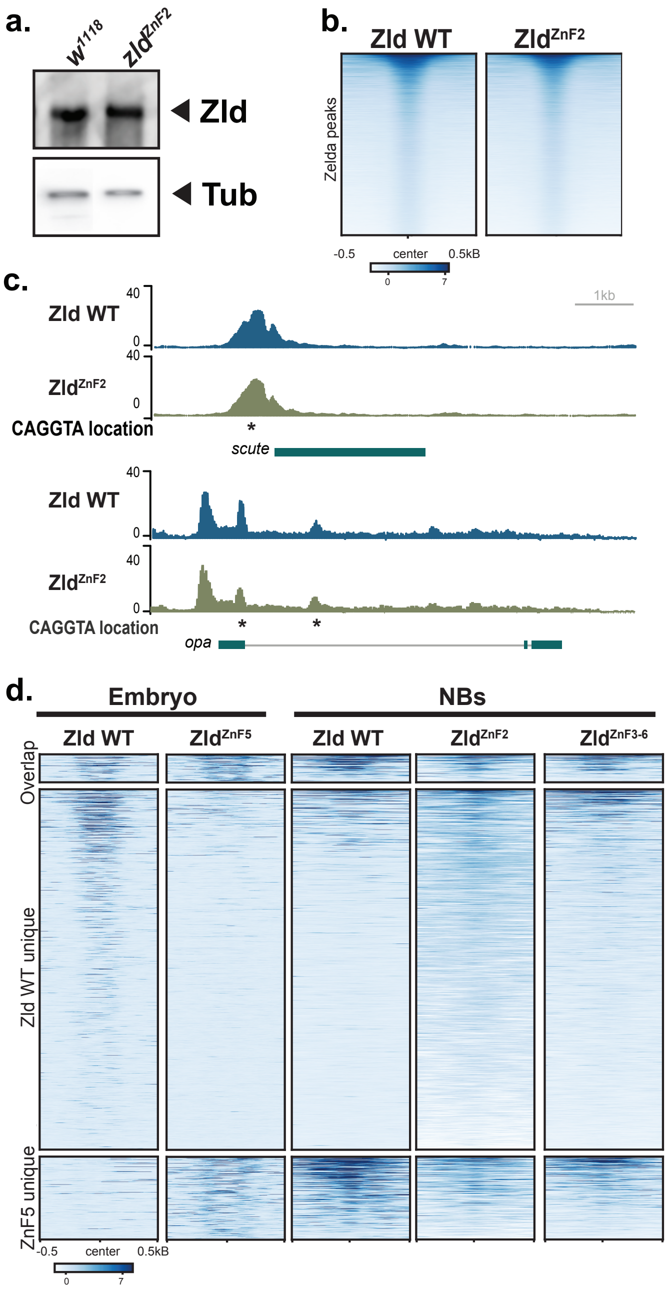


**Supplementary Fig.3: ZnF2 is dispensable for Zld binding in the embryo, but mediates redistribution to regions bound by Zld in neuroblasts. a.** Immunoblot for Zld on stage 5 embryos demonstrate equivalent protein levels for wild type and ZnF2 mutant. Tubulin is shown as a loading control. **b.** Heatmaps of Zld ChIP-seq in wild-type and Zld$^{ZnF2}$ embryos centered on Zld-bound peaks. **c.** Genome browser tracks showing representative loci of up-regulated Zld-target genes from Hamm et al. *PLoS Genet* 2017. Zld binding in wild-type and Zld$^{ZnF2}$ embryos along with CAGGTA motifs are shown. **d.** Heatmaps of Zld occupancy in embryos and neuroblasts for the genotypes indicated above at genomic sites defined by comparing wild-type Zld and Zld$^{ZnF5}$ in the embryo (from Fallacaro et al. bioRxiv 2024).

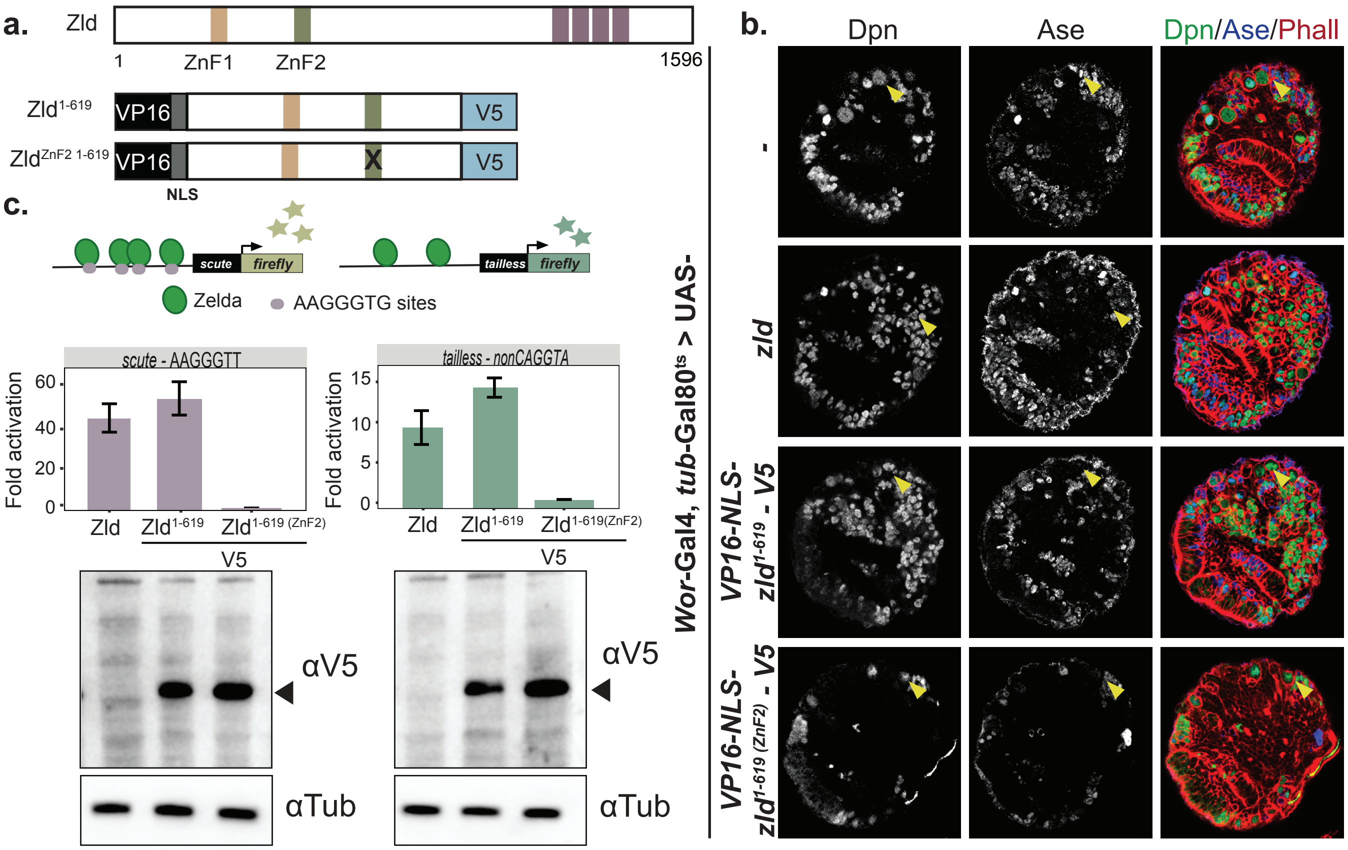


**Supplementary Fig.4: The N-terminus of Zld is sufficient for mediating transcriptional activation. a.** Cartoon schematic of VP16-Zld constructs. **b.** Immunostaining of third instar larval brains lobes expressing ectopic Zld or VP16-NLS constructs as indicated. Expression is driven in type I and type II neuroblasts by *Wor-Gal4, Tub-Gal80$^{ts}$*. Yellow arrows indicate a selection of Dpn+, Ase- type II neuroblasts. Scale bar, 20μm. **c.** Luciferase reporter assays with full-length Zld, Zld$^{1\text{-}619}$ WT and ZnF2 mutant (Zld$^{1\text{-}619\ (ZnF2)}$). Data are shown for both the *scute* reporter with the four CAGGTA replaced with AAGGGT and the neuroblast-specific *tll* enhancer. Immunoblots for V5 tagged proteins indicate protein expression levels. Tubulin is shown as a loading control.
